# Crosstalk between Stromal Cells and Macrophages Shapes Host Immunity to Mycobacteria

**DOI:** 10.64898/2026.02.05.704052

**Authors:** Anne Kathrin Lösslein, Sebastian Baasch, Leonhard Wagner, Kristina Ritter, Florens Lohrmann, Katja Gräwe, Nisreen Ghanem, Jana Neuber, Alexandra Hölscher, Markus Sperandio, Christoph A. Thaiss, Julia Kolter, Christoph Schell, Roman Sankowski, Sagar, Georg Häcker, Christoph Hölscher, Björn Corleis, Philipp Henneke

**Affiliations:** Institute for Infection Prevention and Control, Medical Center - University of Freiburg, Faculty of Medicine, University of Freiburg, Freiburg, Germany; Center for Chronic Immunodeficiency, Medical Center - University of Freiburg, Faculty of Medicine, University of Freiburg, Freiburg, Germany; Arc Institute, Palo Alto, CA, USA; Department of Pathology, Stanford University, Stanford, CA, USA; Infection Immunology, Research Center Borstel, Borstel, Germany; Thematic Translational Unit Tuberculosis, German Center for Infection Research (DZIF), Partner Site Hamburg-Lübeck-Borstel-Riems, Hamburg, Germany; Department of Womeńs and Childreńs Health, Karolinska Institutet, Stockholm, Sweden; Institute of Surgical Pathology, Medical Center - University of Freiburg, Faculty of Medicine, Freiburg, Germany; Faculty of Biology, University of Freiburg, Freiburg, Germany; Walter Brendel Centre of Experimental Medicine, Biomedical Center, Institute of Cardiovascular Physiology and Pathophysiology, Ludwig-Maximilians-Universität München, Planegg-Martinsried, Germany; Institute of Neuropathology, Medical Center - University of Freiburg, Faculty of Medicine, Freiburg, Germany; Clinic for Internal Medicine II, Gastroenterology, Hepatology, Endocrinology and Infectious Disease, Medical Center - University of Freiburg, Faculty of Medicine, Freiburg, Germany; Institute of Medical Microbiology and Hygiene, Medical Center - University of Freiburg, Faculty of Medicine, University of Freiburg, Freiburg, Germany; Institute of Immunology, Friedrich-Loeffler-Institut, Greifswald - Insel Riems, Germany; Centre for Integrative Biological Signalling Studies (CIBSS), University of Freiburg, Freiburg, Germany

**Keywords:** stromal cells, fibroblasts, mycobacteria, macrophages, granuloma, tuberculosis, serous cavities, lung, omentum, cell-cell interactions

## Abstract

Granulomas are disease-defining heterocellular tissue structures in mycobacterial infections. They play a multifaceted role ranging from containing the pathogen to causing tissue destruction. Here, we established a mature peritoneal granuloma model in C57BL/6 mice to investigate the dynamic cell-cell interactions during mycobacterial infection, including long-term immune alterations in serous cavities as important sites of disease manifestation. We found that mycobacteria reside in stromal cells, which actively modulate the local tissue environment and shape macrophage responses, particularly through formation of chemokines and colony-stimulating factor 1. Chronic infection induces sustained reprogramming and diversification of stromal cells toward specialized, immune-like states, including active transfer of mycobacteria to macrophages and a pronounced interferon response. Consequently, stromal cells acquire immunoregulatory properties and support pathogen handling, monocyte recruitment and macrophage maturation, thereby playing a decisive role in granuloma formation and thus in the immune response to mycobacteria.

**HIGHLIGHTS:**

- A novel peritoneal mycobacterial infection model reveals heterocellular crosstalk in mature granulomas.
- Mycobacterial infections persistently reshape immune architecture of serous cavities as important disease sites.
- Stromal cells act as mycobacterial host cells and acquire immune effector functions.
- Stromal cells co-organize the tissue host-pathogen interface by recruiting and directly communicating with bone marrow-derived monocytes.

## INTRODUCTION

Mycobacteria have coevolved with humans over tens of thousands of years, as exemplified by signs of tuberculosis (TB) in approximately 5% of skeletons from a 10,000-year-old settlement^1,2^. *Mycobacterium tuberculosis* (*M.tb*) continues to pose a major global health threat, causing approximately 1.5 million deaths annually^3^. Additionally, the clinical relevance of environmental non-tuberculous mycobacteria (NTM) is increasingly recognized^4^. Their remarkable adaptability enables mycobacteria to survive for years in the host, leading to chronic disease with a wide spectrum of clinical manifestations^5^. Although pulmonary TB is the most common form of the disease, mycobacteria can infect virtually any organ including serous cavities^6^. In the tissue, they induce granulomas, heterocellular structures composed of immune and non-immune cells that evolve over time^7,8^. Granulomas may be viewed as functional containers for mycobacteria, as they isolate the pathogens from the surrounding tissue. However, they also represent potential survival niches and thus sites of disease reactivation, when immune control declines (e.g. during immunosuppression). Furthermore, they are associated with tissue destruction and fibrotic remodeling, both of which impair clearance of infection and reconstitution of tissue homeostasis^9^.

It is estimated that up to 25% of the world’s population has had prior contact with *M.tb*^3^, and approximately 5-10% of individuals with a positive interferon-gamma release assay may reactivate the infection^10^. It is therefore crucial to decipher the dynamic interface between host and pathogen in the tissue, as underlying mechanisms may be exploited for diagnostic and therapeutic purposes^11^. Granulomas mostly consist of macrophages (Mφ), the main target cells for mycobacteria. They are highly dynamic structures, in which Mφ undergo continuous transformation and adaptation processes including epitheloid transformation^12^ and DNA-damage driven multinucleated giant cell (MGC) formation^13,14^, and metabolic changes, in particular lipid droplet formation^15–17^. Already in steady state, Mφ are heterogenous with tissue-specific characteristics and origins^18^. Resident Mφ are embryonically seeded and are, depending on the organ, replenished by bone marrow-derived monocytes. Bacterial infections disturb the immune homeostasis in the tissue, and in case of chronic stimuli lead to long-term alterations of the Mφ compartment^19^. Of note, granuloma formation is also tissue specific. For example in the skin granulomas largely depend on inflammatory monocytes^20^, whereas in the liver resident Mφ (Kupffer cells) form the core of the granuloma^21^. Next to Mφ, granulomas also contain lymphocytes, granulocytes, dendritic cells and NK cells, all of which specifically contribute to the immune defense against mycobacteria^22^. However, in addition to immune cells, non-immune cells, e.g. stromal cells (SC) are structurally and functionally involved in tissue reactions such as abscesses or granulomas. In the inflammatory context, SC influence scar formation and fibrotic transformation. The latter processes also occur during granuloma formation and resolution^9^. Yet, while SC are recognized as a cellular component of granulomas^8^ and although their role in shaping complex immune cell responses is increasingly recognized^23^, their functional role in granuloma formation remains largely unexplored. This is particularly true with respect to their direct interactions with mycobacteria and crosstalk with Mφ. So far, the difficulty in accurately modeling the formation of mature, necrotic granulomas with chronic manifestation in mice complicates the investigation of structural cells in this context^24^. Small numbers of *M.tb* are sufficient to induce granuloma formation with a caseous necrosis in the lung of humans and non-human primates^25^. In contrast, necrotic granuloma formation is hardly achieved in conventional laboratory mouse strains^24,26^, but requires the employment of transgenic or unusual mouse strains^27,28^. Infections of serous cavities, in particular the pleural cavity and to a lesser extent, the peritoneal cavity, are important TB manifestations^29^. Here, we took advantage of the filtering capacity of the omentum, a fold of the peritoneal membrane, to localize the infection and enable spatial analysis of tissue infection dynamics. After intraperitoneal (i.p.) injection of *M. bovis* BCG we found well-organized granuloma structures in the omentum together with chronic inflammation. This novel mature, necrotic granuloma model in C57BL/6 mice enabled detailed characterization of the long-term immune dynamics of granuloma formation, in particular compared to the lung, where mycobacteria-induced infiltrates in C57BL/6 mice are rather diffuse. Using this model, we were able to establish that SC were directly infected by mycobacteria. Infection induced diversification and chronic reprogramming of SC towards immune cell-like behavior and modulation of Mφ recruitment and differentiation. The involvement of SC could be confirmed in single-cell sequencing datasets from human and non-human primate lung granulomas. Thus, cell-intrinsic mechanisms of pathogen handling and induced immunoregulatory properties in SC are key mechanisms in mycobacterial granuloma formation.

## RESULTS

### Intraperitoneal infection with mycobacteria induces necrotic granuloma formation in the omentum accompanied by long-term immune cell alterations

First, we established a model of mycobacterial serous cavities infection as important extrapulmonary disease manifestation. Accordingly, we infected wildtype C57BL/6 mice with *M. bovis* BCG i.p. and focused on events in the peritoneal membrane. We specifically analyzed the omentum, a lipid-rich and mobile part of the peritoneal membrane attached to the stomach, since it functions as a filter for the peritoneal fluid (Figure 1A). The resting omentum is a thin, vascularized clear-white structure. However, 4 weeks post infection (p.i.), we regularly observed the formation of firm tissue knots, often several millimeters in diameter (Figure 1B), although the mice were clinically healthy. Analysis of these structures revealed well-organized granulomas with numerous acid-fast bacilli in the core (Figure 1C, Supplementary Figure 1A). CFU counts confirmed a high bacterial load in the omentum 2 weeks p.i. (Supplementary Figure 1B). The omentum can be considered as a lymphoid organ with lymphoid tissue-like structures forming so-called milky spots^30^. 2 to 4 weeks p.i., when mature granulomas had formed, lymphoid tissue was largely regressed (Supplementary Figure 1C, D). In addition, PAS, AFOG and EvG staining highlighted areas of necrosis at 2 weeks application. The tissue architecture was largely destroyed and nuclear debris was evident (Supplementary Figure 1E). Next to necrosis as a hallmark of mature granuloma formation, epitheloid cells appeared at the boundary zone. 4 weeks p.i. the granuloma size appeared increased and foam cells were detected at higher frequency. At 16 weeks p.i., lymphoid structures and accompanying sclerosis/scarring (i.e. collagen fibers highlighted in blue in the AFOG staining) were evident with only minute amounts of granulocytes (Supplementary Figure 1D, 1E). Granulomas were partially, but not completely, resolved (Figure 1D, Supplementary Figure 1D). Notably, bacteria could no longer be cultured from the omentum at this time point (Supplementary Figure 1B). Moreover, we found granuloma cells to stain positive for γH2Ax, indicating DNA damage (Figure 1E, Supplementary Figure 1F). This was in line with our previous reports that the immune response to mycobacteria is accompanied by DNA damage^13,14^. Overall, these findings showed the formation of mature, necrotic granulomas in the omentum, with an acute phase in the first 2 weeks p.i., and a chronic phase with remaining granuloma structures until at least 16 weeks p.i.. Immunofluorescence staining of mature granulomas at 4 weeks p.i. revealed their heterocellular composition (Figure 1F), with neutrophils and mycobacteria concentrated in the core, which was surrounded by Mφ. The predominance of neutrophils in the granuloma core was supported by flow cytometric analysis, showing a massive increase in neutrophils 24 hours p.i. (Figure 1G). This observation corresponded to mature granuloma formation in humans, in which neutrophils are important contributors^31^. This was followed by incoming monocytes (Figure 1H). To further analyze their recruitment, we infected *Cxcr4^creER^:R26^tdTomato^*mice, in which tamoxifen application stably labels hematopoietic stem cell-derived cells^32^. Recombination was induced 4 weeks before infection (Figure 1I, Supplementary Figure 2A). We found a strong increase in SiglecF^-^ Ly6G^-^ [Lin] Tim4^-^ F4/80^+^ CD11b^+^ both Ly6C^high^ and Ly6C^low^ omental Mφ (Lin^-^ Tim4^-^ F4/80^+^ CD11b^+^) 2 weeks p.i., of which the majority was positive for *Cxcr4*-expression (Figure 1J, Supplementary Figure 2B). Histology indicated a high influx of bone marrow-derived cells to iNOS-positive granulomas (Figure 1K). iNOS is a hallmark of mycobacterial granulomas and a driver of multinucleated giant cell formation^14^ in mice. We detected iNOS-positive cells in the omentum after 16 weeks (Figure 1L), indicating indeed chronic inflammatory changes. Of note, the majority of the iNOS-positive cells coexpressed Mφ markers and in part the inflammatory monocyte marker Ly6C (Figure 1M). Additionally, the cells were tdTomato-positive in *Cxcr4^creER^:R26^tdTomato^*mice showing that recruited cells were the predominant source of iNOS throughout the infection (Figure 1N, O, Supplementary Figure 2C, 2D). This indicated an important role of incoming monocytes for nitric oxide production and mycobacterial containment. In line with long-term immune alterations in the omentum, we detected expression of the programmed death-ligand 1 (PD-L1) by inflammatory monocytes 2 and 4 weeks p.i. in the bloodstream and by CD11b-positive cells in the omentum (Supplementary Figure 2E, 2F). The PD-L1/PD-1 receptor-ligand interaction is widely considered to be immunosuppressive^33^. Human epitheloid granulomas^34^ and Mφ in non-human primate granulomas^35^ were reported to be PD-L1 positive. Thus, this axis might contribute to the chronicity as observed in our model. To explore the role of mycobacterial viability during granuloma formation, we combined fate-mapping with antibiotic treatment (isoniazid plus rifampicin) at 2 weeks p.i. (Figure 1P). The observation that BCG could not be cultured from spleen, liver and bone marrow at 2 weeks p.i. confirmed effectiveness of the antibiotics (Supplementary Figure 2G). However, antibiotic treatment did not alter the Mφ composition of the omentum (Figure 1Q, 1R) or the replacement of Ly6C^+^ and Ly6C^-^ subsets by bone marrow-derived cells after infection (Figure 1R). Moreover, AB treatment did not significantly influence iNOS formation (Figure 1S) or spleen weight (Supplementary Figure 2H), indicating that the chronic immune response was ongoing independent of bacterial viability (Supplementary Figure 1B).

**Figure 1.**
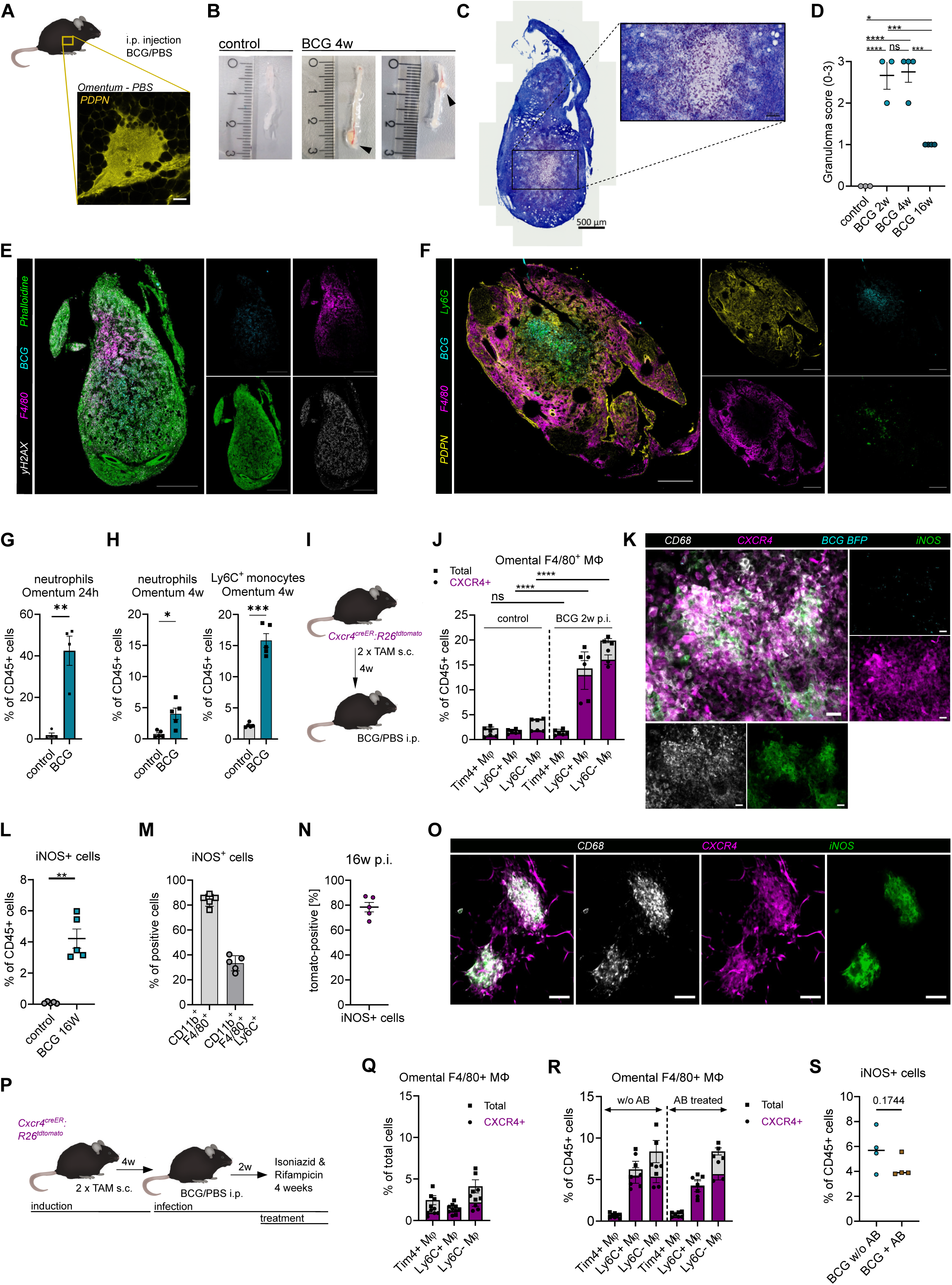
A. Graphical abstract of the intraperitoneal BCG infection model and podoplanin (PDPN) staining of steady state omentum (milky spot). Scale bar 50 µm. B. Omentum of a PBS control mouse (left) and two exemplary BCG i.p. infected mice 4 weeks p.i.. Black arrows highlight macroscopically visible granuloma structures. C. Ziehl-Neelsen staining of a cryo-preserved granuloma isolated from a BCG infected mouse 4 weeks p.i.. Staining was performed on granulomas from 4 mice in independent experiments (compare S1A). D. Depicted is mean ± SEM of granuloma score of omentum from control mice (n=3), BCG mice 2w p.i. (n=3, 1 experiment), 4w p.i. (n=4, 2 experiments), and 16w p.i. (n=4, 1 experiment). Ordinary one-way ANOVA with Tukeýs multiple comparisons test. E. Immunofluorescence staining for F4/80PE, Phalloidin AF488 and (H2AX AF647 of a cryopreserved granuloma 4 weeks p.i.. The granuloma staining was performed in 3 mice independently. Scale bar: 500 µm. F. Immunofluorescence staining for F4/80AF488, Ly6G PE and Podoplanin AF647 of a cryopreserved granuloma 4 weeks p.i.. The granuloma staining was performed in 4 mice independently (maximal intensity projection). Scale bar: 500 µm. G. Depicted is mean ± SEM of neutrophils in omentum 24h p.i. of BCG (n=4) compared to control mice (n=4) in 2 independent experiments (Welch’s t-test). H. Depicted is mean ± SEM of neutrophils and Ly6G^-^ CD11b^+^ Ly6C^+^ cells in omentum and control mice (n=5) and 4w p.i. with BCG (n=5) in 5 independent experiments (Welch’s t-test). I. Graphical abstract of tamoxifen induction of *Cxcr4^creER^:R26^tdTomato^* mice. J. Depicted are means ± SEM of Mφ subsets in the omentum of control (n=3) and BCG-infected *Cxcr4^creER^:R26^tdTomato^* mice (n=3) 2w p.i. with tdTomato-positive percentage displayed in purple (3 independent experiments). 1 mouse was removed, in which insufficient i.p. infection was suspected. Two-way ANOVA; statistical comparisons were performed between PBS and BCG within each subset (all cells and tdTomato-positive cells). K. Immunofluorescence staining for CD68 AF647 and NOS2 AF488 of omentum whole mount of BCG infected *Cxcr4^creER^:R26^tdTomato^* mouse. Staining was performed in at least 4 mice independently. L. Depicted is mean ± SEM of percentage of iNOS-positive CD45^+^ cells in omentum in WT mice (n=5 in 3 independent experiments) M. Surface marker expression of iNOS^+^ cells from 1L. N. Depicted is % of tdTomato-positive iNOS^+^ cells in omentum of *Cxcr4^creER^:R26^tdTomato^* mice 16 weeks p.i. (n=5 in 3 independent experiments) O. Immunofluorescence staining for CD68 AF647 and NOS2 AF488 in omentum whole mount 16 weeks p.i. BCG infection in *Cxcr4^creER^:R26^tdTomato^* mice. The staining was performed in 4 mice in 2 independent experiments. P. Graphical abstract of antibiotic treatment in *Cxcr4^creER^:R26^tdTomato^* mice 2 weeks p.i.. Q. Depicted are means ± SEM of omental macrophage populations (n=5) with and without antibiotic treatment pooled with tdTomato-positive percentage displayed in purple in 3 independent experiments. R. Depicted are means ± SEM of omental macrophage populations of infected mice with antibiotic treatment (n=3-4) and without antibiotic treatment (n=4) in 3 independent experiments. S. Graph shows mean ± SEM of iNOS^+^ cells in omentum from experiment in Fig. 1R (unpaired t-test).

In summary, the mature mycobacterial granuloma model in the omentum of C57BL/6 mice showed granuloma formation for at least 16 weeks, and demonstrated the recruitment of bone marrow-derived monocytes, which were the predominant source of iNOS in the mature granuloma.

### Mycobacterial infection induces long-term alterations in the Mφ compartment of serous cavities

The omentum is surrounded by peritoneal fluid, containing embryonic large (LPM) and monocyte-derived small (SPM) peritoneal Mφ, which are the first immune cells encountering bacteria in infection. When characterizing peritoneal Mφ populations during infection, we observed a long-lasting loss of LPM (CD45^+^, CD115^+^, CD11b^+^, Ly6C^-^, MHC-II^-^, F4/80^+^), whereas the frequency of SPM (CD45^+^, CD115^+^, CD11b^+^, Ly6C^-^, MHC-II^+^, F4/80^-^) increased (Figure 2A, 2B). In flow cytometry, the majority of LPM were BCG BFP positive within 30 minutes (Figure 2C) and downregulated the M-CSF receptor CD115 (Supplementary Figure 3A). After disappearance, they did not repopulate the cavity for several weeks (Figure 2B). The loss of LPM was accompanied by a strong increase in neutrophils in the peritoneal lavage during the first 24 hours (Figure 2D), mirroring the response observed in the omentum (Figure 1G).

**Figure 2.**
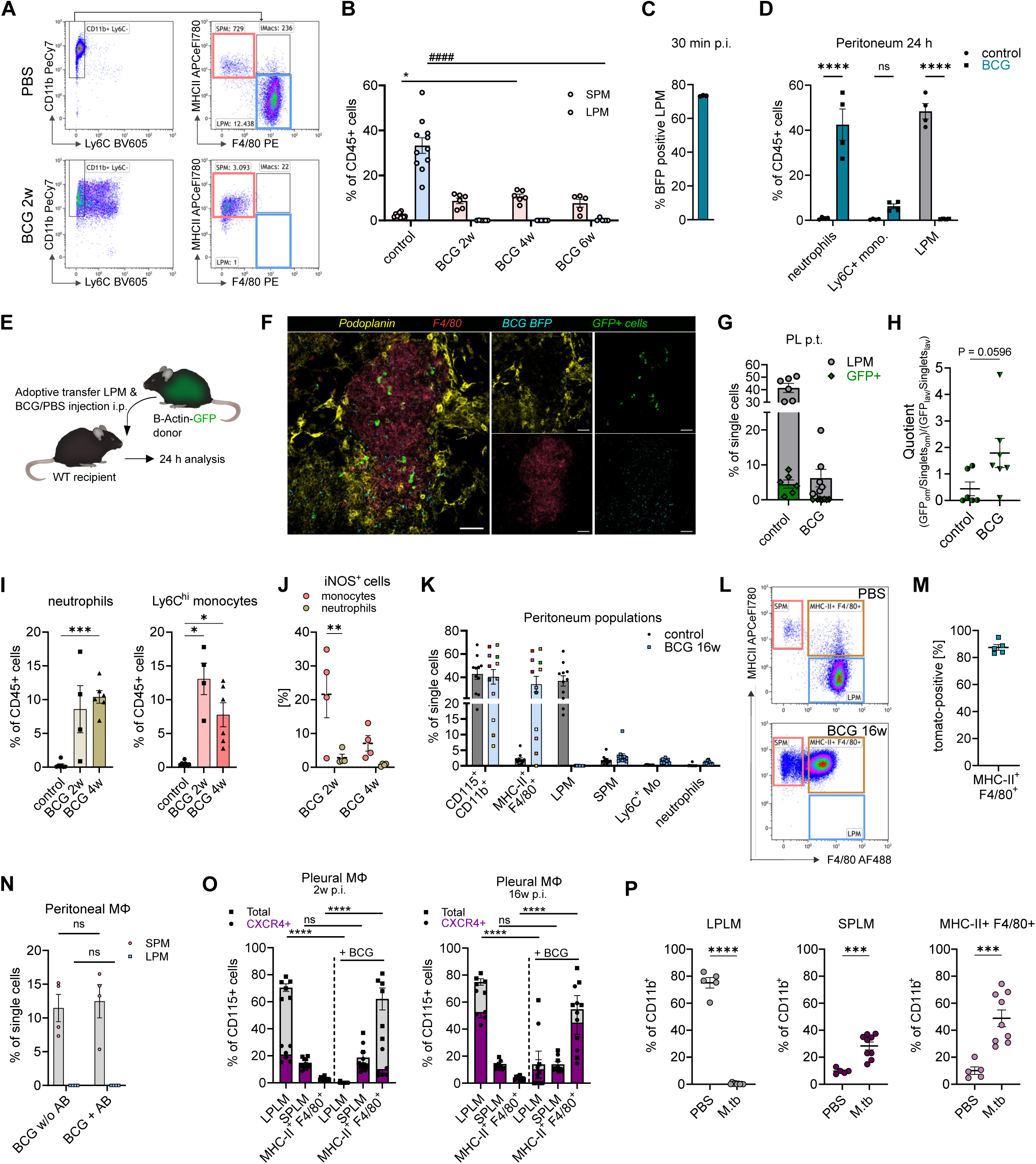
A. Gating strategy depicting differences in macrophage subsets in the peritoneal lavage in control and BCG infected mice. Initial plot was pre-gated on CD115^+^ CD45^+^ single cells. B. Depicted is mean ± SEM of SPM and LPM in control mice (n=11), BCG-infected mice 2 weeks p.i. (n=6), 4 weeks p.i. (n=6), 6 weeks p.i. (n=5) in 10 independent experiments. 2 out of 12 experiments were removed, where insufficient i.p. infection was suspected. Two-way ANOVA with Dunnett’s multiple comparisons test. C. Bar shows mean ± SEM of BFP-positive LPM 30 min post injection of BCG-BFP (n=3 mice of 3 independent experiments). D. Depicted are means ± SEM of neutrophils, Ly6C^+^ monocytes and LPM in PL 24h p.i. of BCG (n=4) compared to control mice (n=4) in 2 independent experiments (two-way ANOVA). E. Graphical abstract of adoptive LPM transfer. F. Immunofluorescence staining for podoplanin AF647 and F4/80 PE of an omentum whole mount of an infected mouse post LPM transfer (compare fig. 2E). The staining was performed in 5 independent mice. G. Depicted is the mean ± SEM of percentage of LPM and transferred GFP^+^ LPM in peritoneal lavage 24h p.t. in PBS (n=6) and BCG-infected mice (n=7) in 7 independent experiments. One PBS mouse was removed due to an unrelated abdominal pathology (hydronephrosis). H. Shown is mean ± SEM of the relation of GFP-positive transferred cells in omentum (om) and PL (lav) relative to single cells (singlets), (unpaired t-test, compare fig. 2E). I. Depicted are means ± SEM of neutrophils and Ly6C^+^ monocytes in PL 2 (n=4) and 4 weeks (n=6) p.i. compared to control (n=7) in 7 independent experiments (Brown-Forsythe and Welch ANOVA). 1 mouse was removed, where insufficient i.p. infection was suspected. J. iNOS staining was performed in 6 experiments of 3I. Depicted is mean ± SEM (Two-way ANOVA). K. Depicted are means ± SEM of immune cell populations in PL of control (n=11) and infected (n=12) WT and *Cxcr4^creER^:R26^tdTomato^* mice 16 weeks post infection (8 independent experiments). L. Exemplary FACS plots of macrophage subsets in PL in control mice and 16 weeks p.i. with BCG. M. Depicted is mean ± SEM of tdTomato-positive cells in the MHC-II^+^ F4/80^+^ population from *Cxcr4^creER^:R26^tdTomato^* mice 16 weeks p.i., if detected (n=5 in 3 experiments). In 1 infected mouse no double positive cells were found. N. Peritoneal macrophage populations with and without antibiotic treatment (n=4 in 3 independent experiments, Two-way ANOVA with Sidak multiple comparisons test). O. Depicted are means ± SEM of small and large pleural macrophages and the percentage of tdTomato-positive cells in *Cxcr4^creER^:R26^tdTomato^* mice 6 weeks post induction for controls (n=6) and 2w-i.p.-infected mice (n=6, in 6 experiments) (left) and mice 20 weeks post induction for controls (n=5) and 16w-i.p.-infected mice (n=6, in 3 experiments (right). The mouse excluded in 1J was also excluded here. P. Graphs show means ± SEM of pleural macrophage populations in control (n=5) and *M.tb* lung infected mice (n=9) in two independent experiments (Welch’s unpaired t-test).

We hypothesized that the loss of Mφ in the fluid was partly due to migration of LPM to the surrounding tissues including the omentum. Therefore, we performed an adoptive transfer of LPM isolated from β-Actin-GFP mice (Figure 2E) and screened for GFP^+^ LPM in omentum whole mounts (Figure 2F). LPM (both total and GFP^+^ LPM) were reduced in the peritoneal fluid of infected mice compared to PBS controls (Figure 2G). Moreover, infection induced a trend towards a relatively increased migration of GFP^+^ cells to the omentum (Figure 2H, Supplementary Figures 3B, C). In addition, we analyzed the percentage of BCG-BFP^+^ immune cells (CD45^+^) 24 h p.i. in the omentum. The majority of the BFP^+^ immune cells expressed the Mφ marker F4/80 (Supplementary Figure 3D). Next to the acute changes in the peritoneal fluid, we analyzed the immune cell composition at later time points of infection. Following the initial influx (Figure 2D), neutrophils decreased 2 weeks p.i. and remained stable (Figure 2I). At 2 and 4 weeks p.i., inflammatory monocytes were significantly increased in the peritoneal lavage (PL) (Figure 2I) and displayed elevated iNOS expression (Figure 2J). 16 weeks p.i., LPM were still absent in the PL, whereas a F4/80 and MHC-II double positive population emerged in most infected mice (Figures 2K, 2L, Supplementary Figure 3E). In *Cxcr4^creER^:R26^tdTomato^*fate mapping mice, about 90% of these F4/80⁺ MHC-II⁺ Mφ were tdTomato-positive 16 weeks p.i. (Figure 2M). Accordingly, the double-positive Mφ originated from hematopoietic/SPM-derived cells rather than LPM-derived cells, which were rarely replaced by monocytes under steady-state conditions (Supplementary Figure 3E). Indeed, LPM may only be replenished by bone marrow-derived cells, namely SPM, in certain conditions, e.g. in C/EBPβ-deficient mice^36^. Antibiotic treatment as described above (Figure 1P) also did not lead to a recovery of the LPM compartment (Figure 2N). The loss of LPM was consistent with a “Mφ disappearance reaction” as described previously for other infection states^37,38^, in which LPM are no longer detectable in the peritoneal fluid due to mechanisms such as migration into surrounding tissues. Next, we wondered, whether the alterations in the peritoneal cavity were representative of other serous cavities. Thus, we analyzed the pleural cavity, which has a similar myeloid composition as the peritoneal fluid under steady-state conditions and is a frequent site of mycobacterial infections^23^. The combination of the i.p. BCG infection with monocyte fate mapping revealed an upregulation of MHC-II rather than a loss of large pleural Mφ upon infection, as the resulting F4/80^+^ MHC-II^+^ pleural Mφ were largely tdTomato-negative (Figure 2O). This indicated a limited monocyte contribution to pleural Mφ 2 weeks p.i. in the intraperitoneal infection model. Nevertheless, in steady state, more than 50% of large pleural Mφ were tdTomato-positive at later time points, indicating a regular replacement by monocytes even without inflammation (Figure 2O). This was in contrast to large peritoneal Mφ (Supplementary Figure 3E). We wondered whether more direct infection of the pleura would induce a long-lasting Mφ loss similar to the peritoneal cavity. Therefore, we explored a pulmonary infection model with *M.tb*. In this model, we indeed observed a long-term loss of large pleural Mφ (LPLM) and an increase in small and MHC-II^+^ F4/80^+^ pleural Mφ comparable to the peritoneal cavity (Figure 2P, Supplementary Figure 3F, Supplementary Figure 3G). In summary, mycobacterial infections not only affected the granuloma tissue, but induced long-term changes in the directly adjacent serous cavities with a remarkable reduction of large serous cavity Mφ, which was not reversible by eradication of bacteria with antibiotics. Moreover, the degree of the Mφ disappearance reaction seemed to depend on the severity and localization of infection.

### Stromal cells serve as mycobacterial host cells and shape the cytokine environment

So far, we found chronic alterations in the immune cell compartments, both at the infection site and in the surrounding cavities. Accordingly, we next explored mechanisms driving these long-term changes. LPM depend on the transcription factor GATA6, induced by retinoic acid (RA). WT1^+^ SC in the omentum provide RA^23^. Therefore, we wondered how SC influenced immune cells and their tissue environment during mycobacterial granuloma formation. Notably, we observed a reorganization of PDPN-positive SC in the omentum already 24 hours p.i. (Figure 2F). The main SC populations in the omentum are CD31^-^ CD45^-^ PDPN^+^ PDGFRA^-^ mesothelial and PDGFRA^+^ fibroblastic cells^23^. Interestingly, we observed a fast (<24 hours) and long-lasting (>16 weeks) shift towards fibroblastic cells after BCG infection (Figure 3A, Supplementary Figure 4A). To unravel the functional role of SC during BCG infection, we infected PDPN-positive SC isolated from the omentum of Life-Act mice with BCG *in vitro* and found bacteria embedded in their actin network (Figure 3B). We observed direct uptake for bacteria with and without antibiotic treatment and fluorescent particles (0.47 µm) when administered to SC in culture (Figure 3C), suggesting active phagocytosis by SC. Moreover, transmission electron microscopy confirmed the intracellular localization of BCG (Figure 3D). To analyze whether ingested mycobacteria could survive within SC similar to Mφ, we determined CFUs from cell lysates after extracellular bacteria in the infected SC culture had been killed with gentamicin 28 hours p.i.. We observed viable bacteria for up to 10 days in culture indicating that SC may serve as potential survival niches (Figure 3E). It has been reported that mycobacteria metabolize host lipids^39^. Moreover, we previously observed that alterations in host cell lipid metabolism are intertwined with the adaptation process of granuloma Mφ subsets^16^. In sorted SC, we found changes in gene expression of *Abca1*, *Dhcr24* and *Fasn*, which are involved in cholesterol and fatty acid metabolism (Supplementary Figure 4B) at different timepoints p.i.. The combination of *Dhcr24* upregulation without *Abca1* upregulation 4 weeks p.i. pointed towards a metabolic shift to cholesterol/steroid accumulation. Additionally, lipid droplet formation was detectable with Nile Red staining after *in vitro* infection (Supplementary Figure 4C). Next, we more deeply assessed the transcriptomic changes in SC after infection by Smart-3 RNA sequencing of isolated SC after 3 days p.i. (Figure 3F, Supplementary Figure 4D). Analysis of differentially expressed genes revealed significantly upregulated GO terms related to bacterial detection and processing, including autophagy and xenophagy in SC after infection (Figure 3G). *Ccl2* and *Ccl7,* which are important for chemotaxis of monocytes, were among the most upregulated genes (Figure 3F and 3H). Importantly, the positive regulation of cytokines, including IL-4, IL-13 and TNF, and nitric oxide biosynthesis were enriched (Supplementary Figure 4E), indicating an active immune effector role for SC. Furthermore, SC infected with BCG showed a significant interferon signature (Figure 3F, Supplementary Figure 4F) and an upregulation of *Gbp2* and *Gbp3* (Figure 3F) encoding Guanylate-Binding Proteins (GBP), which are important in the immune defense against mycobacteria^40^ after infection.

**Figure 3.**
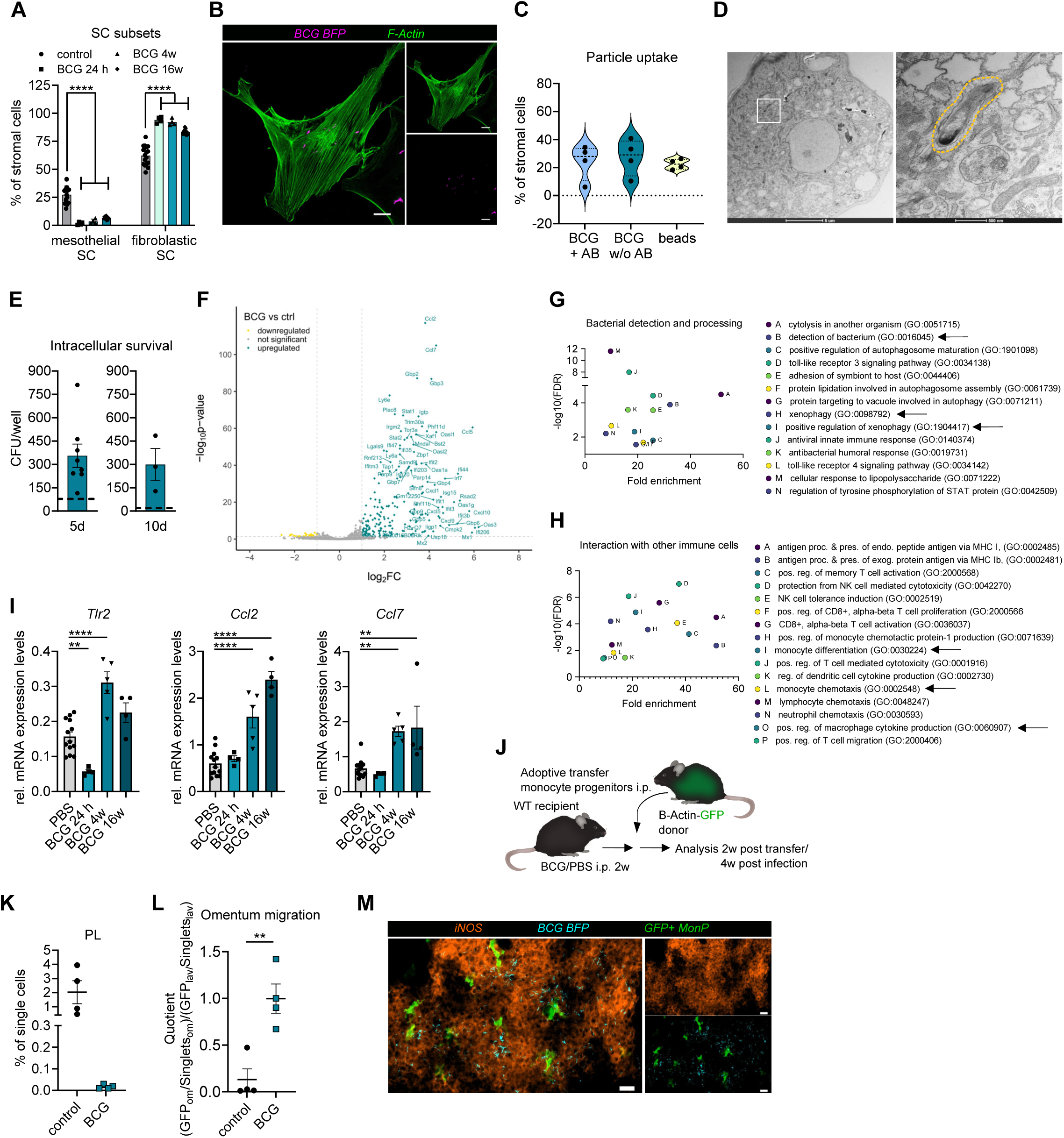
A. Depicted are means ± SEM of mesothelial and fibroblastic stromal cells in omentum in control mice (n=17) and BCG-infected mice at 24h (n=4), 4w (n=4) and 16w (n=9) from WT and *Cxcr4^creER^:R26^tdTomato^* mice in 12 independent experiments (two-way ANOVA with Dunnett multiple comparisons test. B. Imaging of SC culture isolated from Life-Act mice and stimulated for 5 days with BCG-BFP (MOI12) in vitro (n=4 in 2 independent experiments). C. SC were isolated from omentum of WT mice and stimulated with BCG-BFP (MOI12) with and without antibiotics in medium, or biotin-coated fluorescent particles. The percentage of fluorescent SC was evaluated after 3 days (n=4 in 2 independent experiments). D. Electron microscopy of SC infected with BCG (MOI 12), (one experiment, SC pooled from 4 mice). E. SC were isolated from omentum and infected with BCG (MOI12). Gentamicin was added 28h later to kill extracellular bacteria. CFU were plated from SC lysates at day 5 or day 10. The dotted line represents remaining bacterial growth without SC after Gentamicin treatment (n=8 in 5 independent experiments for 5 days, n=3 in one experiment for 10 days). F. Smart-3sequencing of SC cultured with and without BCG (MOI 12), (n=3). Volcano plot depicting differentially regulated genes in infected versus control cells (cutoffs: p ≤ 0.05; │log2FC│ >1; baseMean > 10). G. Depicted are significantly upregulated GO terms in BCG-infected SC related to bacterial detection and processing. H. Depicted are significantly upregulated GO terms in BCG-infected SC related to interaction with other immune cells. I. Relative expression analysis of *Tlr2*, *Ccl2*, *Ccl7* in qPCR of SC directly isolated from omentum of control mice (n=13) and 24h (n=4), 4w (n=5) and 16w (n=4) p.i., (ordinary one-way ANOVA). J. Graphical abstract of monocyte progenitor transfer. K. Depicted is mean ± SEM of GFP positive cells in peritoneal lavage in control (n=4) and infected mice (n=4). Data represent 4 independent experiments. In 1 out of 5 experiments adoptive transfer was unsuccessful in the control and infected mouse (not included). L. Shown is mean ± SEM of the relation of GFP-positive transferred cells in omentum (om) and PL (lav) relative to single cells (singlets), (unpaired t-test, compare fig. 3J). M. Immunofluorescence staining of NOS2 PE and GFP of an omentum whole mount from a BCG infected mouse post adoptive transfer of monocyte progenitors. Staining was performed in 3 mice independently.

To validate these data *in vivo*, we analyzed the expression of *Ccl2* and *Ccl7* in omental SC over the course of infection. Both, *Ccl2* and *Ccl7* increased over time, in line with a constant immune cell recruitment at later time points of infection (Figure 3I). In addition, SC isolated from the omentum expressed *Tlr2* encoding Toll-like receptor (TLR) 2, an important pathogen recognition receptor for mycobacteria, at 4 weeks p.i. (Figure 3I). To further analyze the persistent recruitment of bone marrow-derived monocytes to the infection site, we transferred CD115^+^ CD11b^-^ Ly6C^+^ bone marrow progenitors, including common and induced monocyte progenitors (cMoP, iMoP), into the peritoneum of mice 2 weeks p.i. (Figure 3J). Two weeks post transfer, GFP^+^ progenitor descendants were readily detectable in the peritoneal lavage of control mice, but to a lesser extent in infected mice (Figure 3K, Supplementary Figure 4G). Interestingly, in control mice, progenitor descendants established as a new Mφ state (MHC-II^+^ F4/80^+^ Mφ) (Supplementary Figures 4G, 4H). In contrast, during infection, GFP^+^ cells migrated into the omentum (Figure 3L), in immediate proximity to BCG (Figure 3M). In summary, we found that SC provided a niche for mycobacterial survival and underwent long-term changes during mycobacterial infection, including the acquisition of immune effector functions, thereby promoting monocyte recruitment.

### Stromal cells interact with Mφ and internalize mycobacteria in vivo

Next, we wanted to assess whether SC in the granuloma directly interacted with Mφ, as leading immune cell type in the granuloma. To this end, we established a co-culture model, in which SC were infected for 3 days before the addition of sorted LPM. We found that Mφ and SC closely interacted and that SC released intracellular mycobacteria, which were subsequently taken up by Mφ (Figure 4A, 4B). Importantly, release of the bacteria was not associated with the death of the SC (Figure 4A). The intercellular bacterial exchange as observed by live cell imaging was confirmed in flow cytometry (Figure 4C). This finding revealed a previously unrecognized mechanism of intercellular mycobacterial transfer between non-immune and immune cells with potential implications for bacterial control and containment. Next, we analyzed heterocellular interactions of infected cells *in vivo* (Figure 4D). Adoptively transferred SC, isolated from the omentum of β-Actin-GFP mice, successfully engrafted into the omentum 7 days p.i. (Figure 4E). 3D immunofluorescence analysis showed close interactions of Mφ and SC *in vivo* and pointed towards individual bacteria residing in SC (Figure 4F). To confirm that SC harbored bacteria *in vivo*, we extracted and sorted prior transferred GFP^+^ SC from the omentum of the recipients 10 days post transfer and 3 days p.i. (Figure 4G). Indeed, fluorescence microscopy revealed intracellular bacteria in these cultured SC (Figure 4G, 4H). Next, we explored SC-Mφ interactions in mycobacterial lung infection. Therefore, we adoptively transferred SC isolated from the omentum of control or infected β-Actin-GFP mice into WT recipients via oropharyngeal delivery (Figure 4I). Notably, the transfer of control SC significantly increased the amount of large pleural Mφ in steady state, supporting a regulatory role of SCs in controlling the Mφ composition of serous cavities. Similar to what we observed after *M.tb* lung infection, intratracheal (i.t.) BCG infection triggered a loss of large pleural Mφ, while small and intermediate pleural Mφ expanded (Figure 4L, Supplementary Figure 5A). In infected WT recipient mice, GFP^+^ SC remained detectable 4 weeks after transfer, unlike in non-infected controls (Figure 4J). Despite the small number of transferred SC (Supplementary Figure 5B), immunofluorescence revealed SC in close contact with iNOS–positive Mφ (Figure 4K). In summary, SC phagocytosed mycobacteria and delivered them to Mφ in a controlled manner, indicating a role for SC in barrier function and pathogen containment during infection.

**Figure 4.**
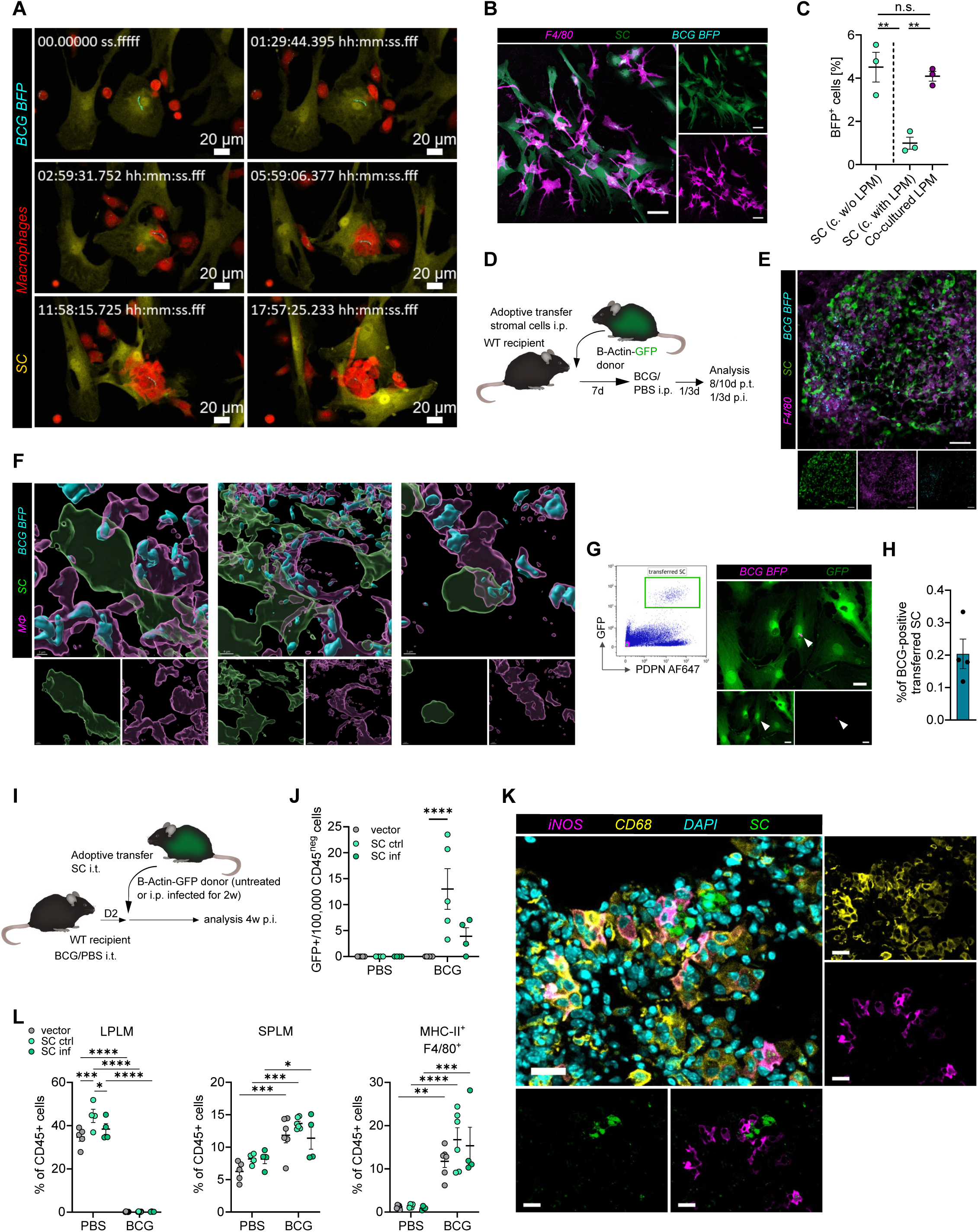
A. Life cell imaging of SC isolated from β-Actin-GFP mice and infected with BCG-BFP. LPM were 3 days later isolated from β-Actin-dsred mice (or vice versa) and added to the SC culture. LCI was started after addition of LPM (3 independent experiments). B. Immunofluorescence staining of co-culture of LPM isolated from WT mice and stained with F4/80 AF488 and SC isolated from β-Actin-dsred mice. Staining was performed in n=4 co-cultures in 2 independent experiments (Scale bar 50 µm). C. SC were infected with BCG and LPM were added on day 3 for cocultures. BCG-BFP positive cells were measured in SC in FACS with and without co-culture with LPM (3 independent experiments, ordinary one-way ANOVA). D. Graphical abstract of adoptive stromal cell transfer. E. Immunofluorescence staining for F4/80 PE and Anti-GFP in infected omentum (1d p.i.) with transferred SC (8d post transfer), (see 4D). The adoptive transfer and staining were performed in n=4 BCG-infected mice in 3 independent experiments. F. 3D reconstruction of immunofluorescence staining in Fig. 4E. G. GFP^+^ SC were sorted from recipients 10d post transfer and 3d post BCG infection (see 4D) and cultured. The gating strategy is represented in the FACS plot. Immunofluorescence staining depicts previously transferred GFP^+^ SC sorted from omentum of recipients. H. Depicted is mean ± SEM of BCG-positive SC from 4G referred to the number of plated SC (n=4 in 3 independent experiments). I. Graphical abstract of the lung infection and oropharyngeal adoptive transfer of stromal cells isolated from the omentum of β-Actin-GFP control or i.p. infected mice. J. Depicted are means ± SEM of GFP^+^ SC referred to 100,000 CD45^-^ cells in control mice with vector (n=5), control SC (n=4) and SC from i.p. infected mice (SCinf, n=4) as well as oropharyngeally infected mice with vector (n=6), steady-state SC (n=5) and SC from i.p. infected mice (SCinf, n=4), (6 independent experiments, two-way ANOVA, Dunnett’s multiple comparisons test). K. Immunofluorescence staining for CD68 AF647, NOS2PE and DAPI of an infected lung post transfer with steady-state SC. L. Shown are means ± SEM of immune cell subsets in pleura lavage. Number of biological replicates depicted in 4J (two-way ANOVA, Tukeýs multiple comparisons test).

### Single-cell profiling uncovers extensive remodeling of both Mφ and stromal compartments in mycobacterial infection

To further characterize changes in SC and Mφ including their functional interactions, we performed single-cell RNA sequencing of F4/80^+^ CD11b^+^ Mφ and CD45^-^ CD31^-^ PDPN^+^ SC 4 weeks p.i.. Clustering analysis revealed that mycobacterial infection induced profound alterations in both Mφ (Figure 5A, Supplementary Figures 6A, 6B) and SC (Figure 5B, Supplementary Figures 6A, 6C), with some clusters being exclusively present in uninfected controls (Figure 5C). The tissue-resident Mφ clusters M3 (expressing *Lyve1* and *Cd163*), M4 (expressing *Cx3cr1*) and M6 (expressing *Timd4* and *Vsig4*) were strongly reduced during infection (Figure 5C–E). Consistent with our previous findings, the infection was dominated by inflammatory monocytes (M5) and activated Mφ with oxidative stress (M0), pro-inflammatory (M1), and interferon (M2) signatures (Figure 5C–E, Supplementary Figure 6D).

**Figure 5.**
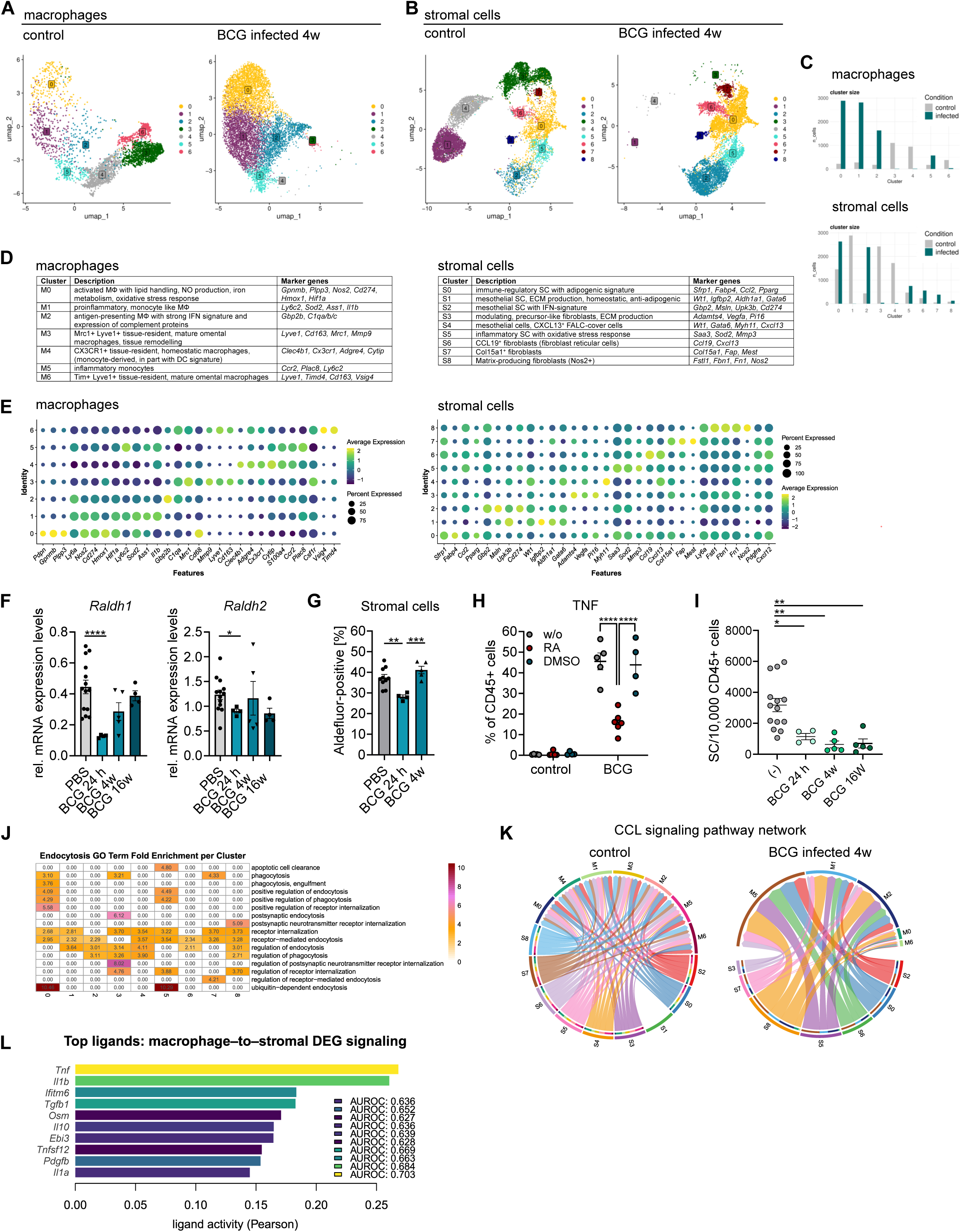
A. F4/80^+^ CD11b^+^ macrophages and PDPN^+^ CD45^-^ CD31^-^ SC were isolated from omentum of control mice (n=5) and BCG infected mice 4w p.i. (n=3) to perform single cell sequencing. Graphs show macrophage clusters in control and infection. B. Graphs show SC clusters from the analysis described in 5A in control and infection. C. Cluster sizes calculated for macrophages and stromal cells. D. Description of macrophage and SC cluster characteristics. E. Depicted are characteristic macrophage and SC markers and their average expression and percent expressed in the individual clusters. F. Relative expression analysis of *Raldh1, Raldh2* in qPCR of SC directly isolated from omentum of control mice (n=13) and 24h (n=4), 4w (n=5) and 16w (n=4) p.i., (Brown-Forsythe and Welch ANOVA with Dunnett T3 multiple comparisons test). G. Depicted are means ± SEM of Aldefluor positive PDPN+ SC compared to DEAB control in PBS mice (n=10) and BCG mice 24h p.i. (n=4) and 4w p.i. (n=5), (ordinary one-way ANOVA with Tukey multiple comparisons test of 7 independent experiments). H. LPM SC Co-Cultures were cultured without (n=5), with retinoic acid (n=5) or DMSO vector control (n=4). Depicted are means ± SEM of TNF expression on LPMs in 3 independent experiments (two-way ANOVA with Tukey multiple comparisons test). I. Depicted is the relation of SC to CD45^+^ cells in control mice (n=14), 24h p.i. (n=4), 4w p.i. (n=5), 16w p.i. (n=5). Ordinary one-way ANOVA of 10 independent experiments with Dunnett multiple comparisons test. J. Analysis of the Fold Enrichment of the GO term endocytosis (GO:0006897) per cluster, including its child GO terms in the stromal cell clusters. K. CellChat interaction analysis of CCL signaling between stromal cell clusters (senders) and macrophage clusters (recipients) in control and infection. L. NicheNet analysis from single cell sequencing dataset comparing the ligand expression on Mφ to differentially expressed genes (receptors) between infected and control SC.

SC clustered into 9 subtypes (Figure 5B, Supplementary Figure 6A, 6C). The *Wt1*-positive mesothelial clusters (S1 and S4) and the precursor-like fibroblastic cluster (S3) were absent during infection (Figure 5B, 5C). The S4 cluster most likely represented *Cxcl13*-positive fat-associated lymphoid cluster (FALC) stromal cells^38^. Cluster S1 showed the lowest *Pdgfrα* expression, aligning with the mesothelial *Pdgfrα*^−^ cluster in steady state^23^. Conversely, we observed increases of cluster S0, an immune-modulatory cluster with an adipogenic signature, and cluster S2, consisting of mesothelial cells with both *Wt1* expression and a prominent IFN signature (Figure 5D, 5E). Smaller populations of inflammatory fibroblasts with oxidative stress response (S5), *Ccl19^+^*fibroblast reticular cells^41^ (S6) and *Col15a1^+^*fibroblasts (S7) also expanded during infection. Comparison of genes upregulated in infected versus control cells in the Smart-3 sequencing (Figure 3) with the scRNA-Seq stromal clusters showed strong overlap, especially with the IFN-rich cluster S2 (Supplementary Figure 6E). GO term analysis of clusters S0, S2, and S5 (Supplementary Figure 6F) revealed enrichment for chemotaxis-related genes (S0), including CCL signaling; high responsiveness to bacterial stimuli, including antigen presentation and IFN responses (S2), and hypoxic responses (S5), reflecting metabolic changes in granulomas (Supplementary Figure 6F) and SC adaptation. These data strongly supported a high adaptability and immune-transition of SC compartment leading to a transcriptional overlap with Mφ. To further specify the SC adaptation in mycobacterial infection, we focused our analysis on i) the metabolic adaptation, ii) the bacterial uptake mechanisms, and iii) the Mφ-SC crosstalk.

We analyzed the enrichment of genes associated with lipid metabolic processes (Supplementary Figure 7A), and observed significant pathway-dependent increases, in particular in S2 and S5 (e.g. cholesterol metabolic process, positive regulation of steroid hormone biosynthetic process). Of interest, the *Wt1*-positive cluster S1 seemed to be mainly involved in the ‘retinoic acid biosynthetic process’ (Supplementary Figure 7A), in line with previous reports^23^. Omental SC are an important source of retinoic acid^23,42^ and thereby can influence the immune landscape in the peritoneal cavity. Notably, under infection cluster S2 seemed to compensate for the absence of cluster S1 by also showing an enrichment for the GO term ‘retinoic acid biosynthetic process’ (Supplementary Figure 7A). Therefore, we analyzed the kinetics of *Raldh1* and *Raldh2* expression, which are key enzymes of the retinoic acid metabolism^42^, together with the Aldefluor enzyme assay of all SC (Figure 5F, 5G). We found a downregulation as early as at 24 hours p.i., in line with the loss of mesothelial cells (Figure 3A), which was restored at 4 weeks p.i., the timepoint of the sequencing analysis. This indicated a potential lack of retinoic acid very early in infection, which other or modified SC then compensated for (e.g. cluster S2). RA is involved in immunoregulatory processes, such as cytokine production. Accordingly, we observed reduced TNF production by Mφ upon addition of RA to the co-culture (Figure 5H). Moreover, the massive influx of immune cells to the omentum reduced the SC-Mφ ratio long-term (Figure 5I). Thus, paracrine signaling effects by metabolites may be attenuated under infection, thereby potentially preventing a homeostatic situation in the omental tissue.

To next examine the potential for bacterial uptake, we analyzed endocytosis-related GO terms (Figure 5J). Positive regulation of endocytosis and phagocytosis, and ubiquitin-mediated endocytosis were enriched in clusters S0 and S5. Ubiquitin-mediated uptake pathways have been reported to be involved in mycobacterial defense mechanisms^43^. Similar to cultured SC (Figure 3G), xenophagy was enriched in cluster S2 and S5, suggesting that these infection-induced subsets are equipped to directly interact with bacteria (Supplementary Figure 7B).

Finally, we used CellChat analysis to investigate Mφ–stromal cell interactions. In both steady state and infection, SC primarily acted as signal senders and Mφ as recipients (Supplementary Figure 7C). Notably, CCL signaling emerged as a critical communication axis, with inflammatory monocytes (cluster M5) serving as key recipients (Figure 5K, Supplementary Figure 7D), confirming our previous findings (Figure 3I). To further characterize how Mφ contribute to the observed SC transformation during infection, we performed a NicheNet analysis (Figure 5L), which predicted *Tnf* as the main Mφ-derived cytokine contributing to the SC phenotype in mycobacterial infection. Of note, the *Tnf*-*Tnfr1* axis was recently described to promote immune-activation in fibroblasts in the skin^44^. Additionally, *Il1b*, *Tgfb1*, and *Oncostatin M*, among others, provided by Mφ are potential mediators influencing the SC phenotype in granuloma formation (Figure 5L).

In summary, specific SC clusters exhibited response patterns similar to Mφ, characterized by interferon signaling and a high metabolic adaptability of SC. Infection-induced alterations in their immune crosstalk, such as reduced RA synthesis and elevated cytokines, may modulate the immune cell landscape in mycobacteria infection.

### Stromal cells shape the immune cell composition of granulomas

Our data so far suggested that SC were essential for the recruitment of monocytes to the site of mycobacterial infection, thereby potentially impacting heterocellular granuloma development. Accordingly, we wondered if a reduction of SC affected granuloma composition and monocyte recruitment. We employed *Pdgfrα^creER^:R26^DTA/+^* mice, in which SC were reduced by tamoxifen treatment for 5 days starting at day 2 p.i., when most omental SC already expressed PDGFRA (Figure 6A). Under these conditions, the absolute number of PDGFRA^+^ SC was decreased at 4 weeks p.i., i.e. at the peak of granuloma formation (Figure 6B). In addition, the weight of the omentum (Figure 6C) and CD11b^+^ F4/80^+^ omental Mφ were reduced in DTA mice (Figure 6D, Supplementary Figure 8A). We observed strong reductions in absolute cell counts for resident Tim4^+^ Mφ (Figure 6E, Supplementary Figure 8A), double-negative Mφ (Figure 6F, Supplementary Figure 8A), and Ly6C^+^ incoming Mφ (Figure 6G, Supplementary Figure 8A), although the percentage of CD45^+^ cells and frequency of iNOS-positive Mφ remained unchanged (Supplementary Figure 8B, 8C). These data indicated that alterations of the SC niche also had an influence on monocyte recruitment and Mφ composition at the infection site. Moreover, spleen weight was significantly reduced – potentially indicating lower overall inflammation while mice showed less weight and fewer PDGFRA^+^ SC (Supplementary Figure 8D), suggesting potential systemic effects of the depletion.

**Figure 6.**
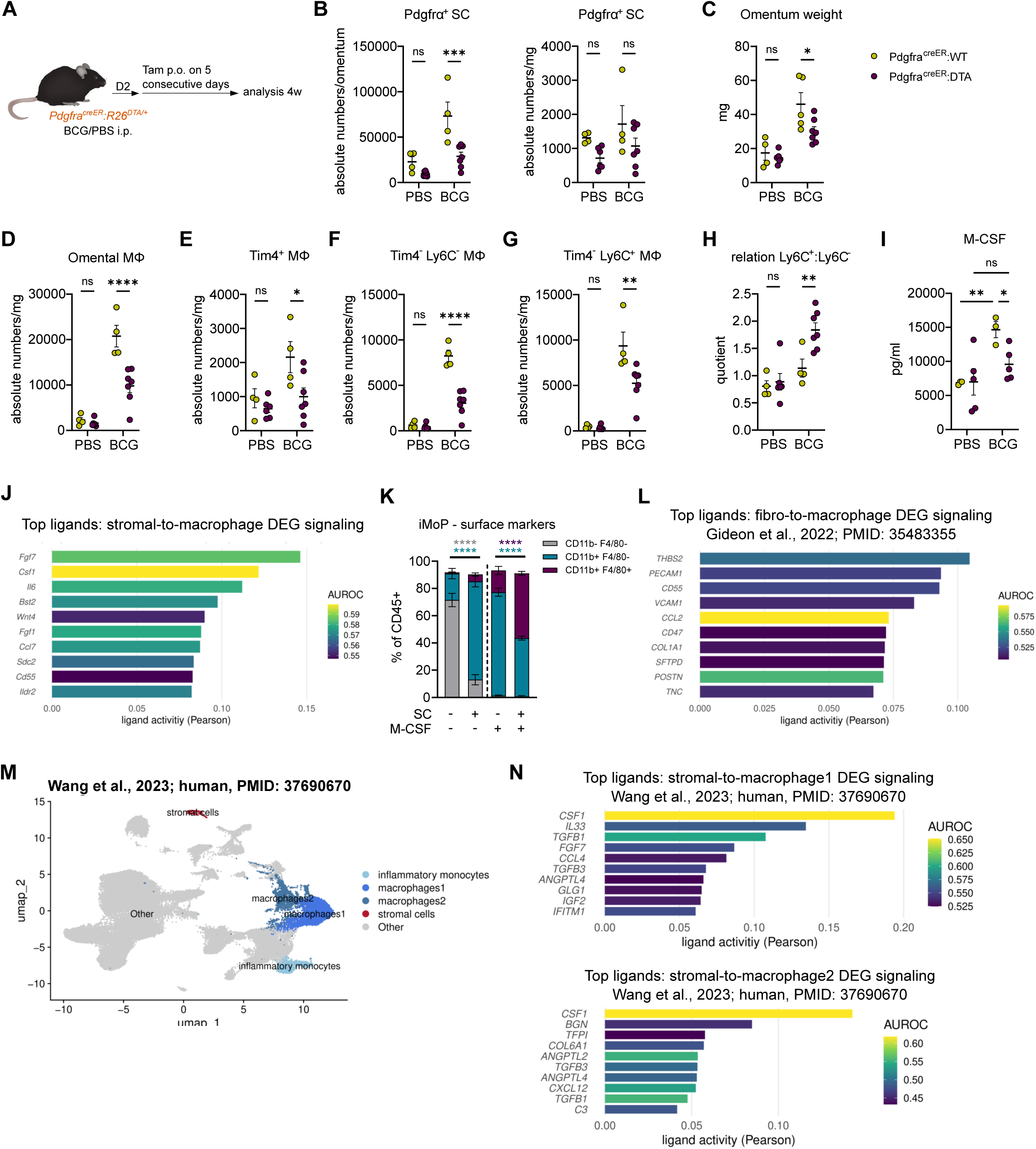
A. Graphical abstract of SC reduction in *Pdgfrα^creER^:R26^DTA/+^* mice with tamoxifen. B. Depicted are means ± SEM of absolute numbers of PDGFRA^+^ SC per omentum and per mg in *Pdgfrα^creER^:R26^+/+^* control (n=4) and infected mice (n=4) and *Pdgfrα^creER^:R26^DTA/+^* control (n=6) and infected mice (n=7) in 5 independent experiments (two-way ANOVA with Sidak multiple comparisons test). One *Pdgfrα^creER^:R26^+/+^* mouse with insufficient infection (no CFU counts) was removed (only stated once for complete fig. 6). C. Depicted are mean ± SEM of omentum weights in *Pdgfrα^creER^:R26^+/+^* control (n=4) and infected mice (n=5) and *Pdgfrα^creER^:R26^DTA/+^* control (n=6) and infected mice (n=7) in 5 independent experiments (two-way ANOVA with Sidak multiple comparisons test). D. Shown are means ± SEM of absolute omental macrophages numbers per mg, (two-way ANOVA with Sidak multiple comparisons test), (compare 6B) E. Shown are means ± SEM of Tim4^+^ macrophages in omentum per mg (two-way ANOVA with Sidak multiple comparisons test), (compare 6B). F. Shown are means ± SEM of Tim4^-^ Ly6C^-^ macrophages in omentum per mg (two-way ANOVA with Sidak multiple comparisons test), (compare 6B). G. Shown are means ± SEM of Tim4^-^ Ly6C^+^ macrophages in omentum per mg (two-way ANOVA with Sidak multiple comparisons test), (compare 6B). H. Depicted are means ± SEM of the relation of Ly6C^+^ Tim4^-^ and Ly6C^-^ Tim4^-^macrophage subsets in omentum (compare 6B). I. Depicted are means ± SEM of M-CSF serum levels. J. NicheNet analysis from single cell sequencing dataset (fig. 6) comparing the ligand expression on stromal cells to differentially expressed genes (receptors) between infected and control macrophages. K. In vitro co-culture of iMoP isolated from bone marrow of WT mice +/- stromal cells isolated from omentum, +/- M-CSF. Surface marker expression was analyzed by FACS 24 hours after plating. Depicted is mean ± SEM of 4-5 replicates in 2 independent experiments (two-way ANOVA with Sidak multiple comparisons test between conditions +/-SC). L. NicheNet analysis from a published single cell sequencing dataset of macaques^9^ comparing the ligand expression on fibroblasts to differentially expressed genes (receptors) between macrophages in high and low burden granulomas. M. Clustering and annotation of macrophages, inflammatory monocytes and stromal cells in a published human single cell sequencing data set of granulomas with high and low burden^45^. N. NicheNet analysis from fig. 6M comparing the ligand expression on stromal cells to differentially expressed genes (receptors) between macrophages of high and low burden granulomas.

Interestingly, SPM were increased in the peritoneal lavage (Supplementary Figure 8E), indicating that the replenishment of the peritoneal compartment was not affected (Supplementary Figure 8E). In contrast, ablation of SC with consequent reduction of granuloma Mφ did not affect bacterial burden at 4 weeks p.i., indicating that ongoing monocyte recruitment after the initial phase might not substantially contribute to bacterial clearance (Supplementary Figure 8F). These results aligned with our observations following antibiotic treatment, wherein immune processes within the granuloma became dissociated from bacterial viability (Figure 1). We also noted a change in the Mφ composition, with the ratio of Ly6C^+^ to Ly6C^−^ cells increasing toward the Ly6C^+^ subset in *Pdgfrα^creER^:R26^DTA/+^*mice, which likely represents the more immature subset (Figure 6H). In line with these findings, we measured reduced concentrations of Mφ colony-stimulating factor (M-CSF) in the circulation of *Pdgfrα^creER^:R26^DTA/+^* mice compared to the *Pdgfrα^creER^:R26^+/+^* controls (Figure 6I). Furthermore, we identified M-CSF signaling as important ligand-receptor interaction in the NicheNet analysis (Figure 6J). We next performed a co-culture of SC and bone marrow-sorted induced monocyte progenitors (iMoP)^16^. We observed that in steady state iMoP upregulated both CD11b and F4/80 faster when cultured together with SC (Figure 6K) in the absence of M-CSF, indicating that the presence of SC can promote Mφ maturation. Finally, we wondered whether SC played a similar role in monocyte recruitment and Mφ maturation in humans and non-human primates. First, we performed a NicheNet analysis on a 10-week scRNA-Seq set by Gideon et al.^9^, analyzing high and low burden granulomas in Macaques. We found *CCL2* as a top ligand when analyzing the influence of fibroblasts on differentially expressed genes between Mφ in high versus low burden granulomas (Figure 6L). These data corresponded to our findings, that fibroblasts produce CCL and thus promote monocyte recruitment even at late timepoints (Figure 3I). In a human data set by Wang et al.^45^, we focused on the interaction of SC with Mφ and inflammatory monocytes (Figure 6M). Comparing high versus low burden lesions, we found both *CSF1* (encoding for M-CSF) and *CCL4* as top hits for genes of soluble ligands in the NicheNet interaction analysis (Figure 6N, Supplementary Figure 8G).

Altogether, our findings in mice, non-human primates and humans demonstrated that SC critically influence the granuloma Mφ composition in mycobacterial infections as well as the recruitment of inflammatory monocytes and their maturation into Mφ.

## DISCUSSION

Granuloma formation in mycobacterial infection is a highly dynamic process in which immune cells adapting to the infectious niche are intertwined with continuous tissue reorganization^46^. Here, we developed a model of serous cavity mycobacterial infection that induces formation of mature, necrotic granulomas in the omentum, thereby revealing a previously underappreciated contribution of stromal cell programming and diversification to the in-tissue host-pathogen interface. The focal establishment of mycobacterial infections followed kinetic and spatial trajectories defined by multiple switch points of heterocellular interactions. After bacterial uptake by LPM within minutes, the immune cell wave in the first 24 hours was dominated by neutrophils. On the one hand this response may represent a rather general reaction to bacterial infection; on the other hand, in the case of mycobacteria, it seemed to pave the way for the formation of the necrotic granuloma core. This is in line with reports that neutrophil NETosis can be associated with granuloma caseation and necrosis^47^. In contrast an early, i.e. potentially exaggerated, neutrophil response may hinder resolution of infection as recently shown by Gern et al.^48^. In the serous cavity model, we found recruited neutrophils to be increasingly replaced by monocytes within the first 2 weeks p.i.. Remarkably, inactivation of bacteria by antibiotics did not alter Mφ composition, suggesting that immune cell recruitment may follow a self-sustaining program once initiated, rather than being acutely dependent on bacterial viability thus constitute a safety layer; however such limited regulation may also promote paradoxical reactions following TB treatment which occur for example when immunosuppression ceases (e.g. in HIV treatment) and which must be differentiated from disease relapse^49^.

Interestingly, the cellular response to mycobacteria was not restricted to the infected tissue. The almost complete loss of the embryonically seeded large peritoneal Mφ was consistent with a Mφ disappearance reaction, a phenomenon described in response to strong inflammatory stimuli^50^. LPM engulfed mycobacteria within minutes, similar to previous reports for *E. coli*^51^, and then relocated to surrounding tissues including the omentum. Even 16 weeks p.i., the regular MHC-II^-^ F4/80^+^ LPM population remained absent from the peritoneal cavity and the myeloid niche was regularly, but not uniformly, filled by monocyte-derived MHC-II^+^ F4/80^+^ Mφ. It seems notable that LPM loss is not specific for mycobacteria since it is observed after high-dose i.p. treatment with zymosan in a sterile peritonitis model^19^. However, Louwe et al.^19^ observed replacement by bone marrow–derived MHC-II⁺ F4/80⁺ cells as early as 3 weeks post-treatment. In contrast, our 16-week time point indicates a markedly prolonged disruption of peritoneal homeostasis during chronic bacterial infection. In addition, even 4 weeks after antibiotic treatment, immune cell composition continued to be disturbed both in the peritoneal cavity and in the omentum. The sustained production of nitric oxide synthase and the disruption of resident Mφ in the omentum may contribute to long-term alterations in the peritoneal cavity, as the omentum plays a key role in maintaining immune homeostasis in the serous fluid^52^. Large peritoneal Mφ are dependent on GATA6^53^ and retinoic acid^23^, which are in part derived from omental SC^23^. It has previously been reported, that the serous cavities, i.e. peritoneum, pleura and pericardium, share a similar Mφ composition in steady state^23,54^ and that pleural Mφ can translocate to the lung tissue in influenza infection^55^. Indeed, we observed a long-term loss of large pleural Mφ with an increase of a double positive MHC-II^+^ F4/80^+^ Mφ population in mycobacterial infection. Yet, despite the similar reaction patterns, the replacement kinetics of large pleural and peritoneal Mφ differed in steady state with only large pleural Mφ being replaced by bone marrow-derived cells. This points towards specific adaptations to tissue-specific requirements, since acute pulmonary infections occur frequently throughout life, whereas the peritoneal cavity is programmed to sporadic microbial translocation from the gut. Mφ are considered to be the key targets for mycobacteria, leading to reprogramming of the Mφ differentiation^13^ and metabolism^16,56,57^. Mycobacteria can manipulate the host cell^7,58^ and benefit from host’s lipids for their own survival^39^. Finding suitable intracellular niches, allows the mycobacteria to persist in the host for ages in dormant states^59–61^ with potential reactivation in case of immunosuppression. However, mycobacteria are in part promiscuous microorganisms that can infect non-immune cells, e.g. glial cells, such as Schwann cells by *M. leprae*^62^. In addition, fibroblasts have been recently shown to become infected by mycobacteria *in vitro* and to alter their cytokine responses upon infection^63^. We observed that mycobacteria directly infected SC both *in vitro* and *in vivo*, supporting the concept that non-immune cells can serve as long-term pathogen niches in chronic infection. Additionally, PDGFRA^-^ mesothelial SC were consistently reduced compared to PDGFRA^+^ fibroblastic SC indicating rapid, long-lasting alterations in SC induced by mycobacterial infection. SC intimately interacted with Mφ, with bacteria being handed over from stromal cells to Mφ, strongly indicating an immune regulatory function of SC. Of interest, SC appeared to survive bacterial release, resembling non-lytic expulsion such as vomocytosis^64,65^. However, vomocytosis has been primarily reported in professional phagocytes, particularly in the context of fungal infections^66,67^. This novel heterocellular interaction mechanisms may have important implications for bacterial control during infection. Accordingly, previous studies indicate that non-lytic expulsion processes may also play a role in bacterial infections, including mycobacterial infections demonstrated in a *Mycobacterium marinum* - amoeba model^68^.

In addition, we found that SC underwent substantial diversification during infection, with specific clusters displaying interferon-signatures, metabolic adaptations and bacterial uptake capacity, highlighting their pronounced plasticity in inflammatory environments. These findings are in line with reports that SC, in addition to their barrier role, also possess immunoregulatory functions^69,70^. Granuloma Mφ appear to contribute to the inflammatory SC phenotype primarily through TNF ligand–receptor interactions with additional contributions from IL1β and OSM consistent with reports from other inflammatory settings^44, 71^. OSM has been shown to activate fibroblasts and promote an inflammatory SC phenotype in tumor^72^ and inflammatory^73^ models. However, these effects appear to be model- and context-dependent, as suppressive roles have also been reported in the heart^74^. Moreover, SC showed a high degree of adaptability and plasticity. To demonstrate, that SC are important for the immune recruitment to the site of infection, we used *Pdgfrα^creER^:R26^DTA/+^* mice^75^. It should be noted that this mouse model does not lead to complete SC depletion and that depletion is not restricted to the omentum, i.e. PDGFRA^+^ SC in other tissue compartments may also be affected. Whereas Mφ and monocyte subsets were reduced in absolute numbers, mycobacterial counts remained unaffected. This observation raised the possibility that myeloid recruitment beyond a certain threshold may no longer promote bacterial clearance but instead reflect a dissociated immune process, potentially increasing immunopathology. Apparently, SC significantly contributed to this rather exaggerated Mφ response. Ongoing monocyte recruitment orchestrated by SC may establish an immune feedback circuit, potentially amplifying immune cell influx independent of bacterial burden. These considerations are consistent with an ambivalent role of the granuloma, balancing pathogen control and an excessive immune response. Beyond recruitment, we identified SC as important source of M-CSF, thereby influencing Mφ differentiation. This observation is in line with reports on SC as source for M-CSF in other diseases and even in homeostasis^76,77^. In particular, our data indicate that SC-derived M-CSF contributes to Mφ-differentiation within the granuloma niche, thereby supporting structural consolidation rather than directly enhancing antimicrobial activity. Granuloma formation has been well described to affect the omentum in humans^78^, although the lung remains the primary target of TB. Importantly our analyses of non-human primate^9^ and human^45^ granulomas revealed conserved stromal-myeloid signaling pathways, supporting the translational relevance of our findings in the omental mouse model.

SC are found throughout all organs and increasing evidence suggests that they have, depending on the subtype, immunoregulatory functions and can adapt to chronic inflammation^79,80^. SC have previously been reported to be part of granuloma structures, for example in leprosy^81^. A recently published sequencing study of human granulomas suggested that fibroblasts act as important signaling cells in cell–cell communication^82^. Likewise, mesenchymal stem cells, as non-immune cells, are gaining increasing attention in tuberculosis research as potential therapeutic targets and as possible host cells supporting bacterial persistence^83,84^.

Together, our results position SC as an active, immunomodulatory component of the granuloma. Its interactions with Mφ and monocytes may represent promising targets for host-directed therapeutic strategies in mycobacterial infections. Thus, our study provides an important foundation for understanding the influence of SC on immune cells and granuloma formation in mycobacterial infections, as well as Mφ-SC interactions in general.

## ACKNOWLEDGMENT

We thank Adriana Greco and Reem Al-Sumati for excellent technical assistance. We are grateful for expert assistance in cellular analysis and imaging to Jan Bodinek-Wersing and the Lighthouse Core Facility, which is funded in part by the Medical Faculty, University of Freiburg (Project Numbers 2023/A2-Fol; 2021/B3-Fol), the DKTK, the Mertelsmann Foundation, and the DFG (Project Number 450392965). Furthermore, we thank the Center for Experimental Models and Transgenic Services (Freiburg) and the animal facilities in Borstel. Additionally, we thank the Core Facility for Electron Microscopy (EMcore) at the University Freiburg Medical Center—IMITATE is registered with the DFG under the reference number RI_00555. We thank Björn Zessin (Institute of Immunology, Friedrich-Loeffler-Institut, Greifswald) for assistance with R scripts. AKL received an individual research grant (2024) from the European Society of Clinical Microbiology and Infectious Diseases (ESCMID). AKL and FL were supported by the clinician scientist program IMM-PACT, funded by the Deutsche Forschungsgemeinschaft (DFG; German Research Foundation) – Project ID 413517907. AKL received funding from the German Research Foundation (DFG, TRR167 [Project ID 259373024]). KR and CH are supported by the German Federal Ministry of Research, Technology and Space (BMFTR) through the German Center of Infection Research (DZIF; TTU-TB, grant number 02.717). The authors acknowledge the support of the Freiburg Galaxy Team: Björn Grüning, Bioinformatics, University of Freiburg (Germany), funded by the German Federal Ministry of Education and Research BMFTR grant 031 A538A de.NBI-RBC and the Ministry of Science, Research and the Arts Baden-Württemberg (MWK) within the framework of LIBIS/de.NBI Freiburg.

RS is supported by the IMMediate Advanced Clinician Scientist-Program, Department of Medicine II, Medical Center, University of Freiburg and Faculty of Medicine, University of Freiburg, funded by the Bundesministerium für Bildung und Forschung (Federal Ministry of Education and Research), 01EO2103. Furthermore, RS is supported by the Fritz Thyssen Foundation. PH received funding from the German Research Foundation (DFG, HE3127/12, HE3127/16, TRR359 [Project ID 491676693], TRR167 [Project ID 259373024], and CRC1160 [Project ID 256073931]).

## AUTHOR CONTRIBUTION

AKL designed the project and performed the majority of the experiments. AKL and PH led the study and wrote the first draft of the manuscript. GH, JK, MS, CAT, SB provided critical intellectual input on study design, methodology and manuscript revision. SB helped with oropharyngeal infections. LW assisted with performing confocal imaging and performed the LEGENDplex™. NG performed electron microscopy and JN prepared the BCG stocks and performed the 3D analysis. FL supported AKL with the scRNA-Seq analysis. S performed the scRNA-Seq and helped with the analysis. KR, AH and CH performed *M.tb* lung infection experiments. KG performed the immunohistochemistry of lungs and histology stainings of omentum. CS evaluated necrosis, quantified lymphoid and granuloma scores of omentum. RS performed Smart-3 sequencing. BC provided R scripts for the reanalysis of published single cell sequencing data. All authors edited the manuscript.

## DECLARATION OF INTERESTS

The authors declare no competing interests.

## DECLARATION OF GENERATIVE AI TOOLS AND AI-ASSISTED TECHNOLOGIES

AI tools were employed to assist in generating initial R scripts. All R scripts were controlled and revised several times by the authors.

## FIGURE LEGENDS

**Supplementary Figure 1**

A. Ziehl-Neelsen staining of cryo-preserved granulomas isolated from BCG infected mice 4w p.i. (compare Fig. 1C).
B. CFU counts in omentum 2 and 16 weeks p.i.. Depicted is mean ± SEM of log counts; n.d. = not detected.
C. Depicted is mean ± SEM of lymphoid scores of omentum from control mice (n=3), BCG mice 2 weeks p.i. (n=3, 1 experiment), 4w p.i. (n=4, 2 experiments), and 16w p.i. (n=4, 1 experiment). Ordinary one-way ANOVA with Tukeýs multiple comparisons test.
D. Representative H&E staining of Fig. 1D/Supp. Fig. 1C. Dashed boxes indicate areas of increased magnification.
E. Representative PAS (Periodic Acid-Schiff), AFOG (Acid Fuchsin-Orange G), EvG (Elastica-van-Gieson) staining 2 and 16 w p.i. with BCG (dashed boxes indicate areas of zoom-in). n=3 animals per time point.
F. Close-up of immunofluorescence staining in Fig. 1E.

**Supplementary Figure 2**

A. Depicted are means ± SEM of percentage of tdTomato-positive Ly6C^+^ monocytes in the bloodstream 6- and 20-weeks post induction in control and infected mice (2w/16w p.i.); n=6 for 2w, n=5-6 for 16w.
B. Gating strategy for omentum populations in *Cxcr4^creER^:R26^tdTomato^* mice.
C. Immunofluorescence staining for CD68 AF647 and NOS2 AF488 of omentum whole mount of a *Cxcr4^creER^:R26^tdTomato^* control mouse 2 weeks after PBS i.p. injection.
D. Gating strategy for iNOS-positive/tdTomato-positive cells in omentum of *Cxcr4^creER^:R26^tdTomato^* mice 16 weeks p.i..
E. Gating strategy for PD-L1 positive inflammatory monocytes in the bloodstream.
F. Depicted are means ± SEM of the % of PD-L1 positive inflammatory monocytes in the bloodstream (left) of control mice (n=6), and BCG-infected mice 2w (n=3) and 4w (n=4), (4 independent experiments in total, Ordinary one-way ANOVA with Dunnett’s multiple comparisons test). On the right, means ± SEM of PD-L1^+^ CD11b^+^ immune cells in omentum are depicted in control (n=4) and infected mice (n=4) in 4 independent experiments (unpaired t-test).
G. CFU counts in spleen, liver and BM post antibiotic treatment.
H. Shown is mean ± SEM of spleen weight with and without antibiotic treatment.

**Supplementary Figure 3**

A. MFI of CD115 expression of LPM 30 min post injection of BCG relative to control mice. Depicted is mean ± SEM of 3 independent mice in 3 experiments (one sample t-test). FACS dot plots shows BFP-positivity and CD115 expression of infected (blue) mice and controls (red).
B. Gating strategy of transferred LPM in omentum.
C. Gating strategy of transferred LPM in peritoneal lavage.
D. FACS analysis of BFP-positive CD45^+^ cells (left) and percentage of F4/80 expression on these CD45^+^ BFP^+^ cells (right) in the omentum 24h p.i. from the BCG-infected mice in 2G.
E. Depicted are means ± SEM of tdTomato-positivity of SPM and LPM 6w post tamoxifen in control (n=6) and infected mice (2w, n=6) and 20w post tamoxifen in control (n=5) and infected mice (16w, n=6).
F. Gating strategy of pleural macrophages in control and *M.tb* infection.
G. Summary of the populations in 2P.

**Supplementary Figure 4**

A. Gating strategy of mesothelial and fibroblastic SC in omentum.
B. Relative expression analysis of *Abca1, Dhcr24, Fasn* in qPCR of SC directly isolated from omentum of control mice (n=8-9), 4w (n=4-5) and 16w (n=4) p.i., (Ordinary one-way ANOVA with Dunnett multiple comparisons test).
C. Immunofluorescence staining for NileRed, F4/80 AF488, PDPN AF647 of SC and LPM co-cultures (n=4 in 3 independent experiments).
D. PCA plot of sequencing described in 3F.
E. Depicted are significantly upregulated GO terms in BCG-infected SC related to cytokines and chemokines.
F. Depicted are significantly upregulated GO terms in BCG-infected SC related to interferon.
G. Gating strategies for adoptively transferred monocyte progenitors in peritoneal lavage and omentum.
H. Distribution of GFP^+^ CD115^+^ transferred progenitors in the respective gates in peritoneal lavage.

**Supplementary Figure 5**

A. Gating strategy of pleural macrophages in control and BCG infection.
B. Depicted are absolute numbers of GFP positive cells from the experiment in 4I/4J.

**Supplementary Figure 6**

A. Complete macrophage and stromal cell clusters (control and infected pooled).
B. Heatmap of the top 10 upregulated genes (min.pct 0.25, logthreshold 0.25) in each macrophage cluster.
C. Heatmap of the top 8 upregulated genes (min.pct 0.25, logthreshold 0.25) in each stromal cell cluster.
D. GO Enrichment analysis for upregulated genes in macrophage clusters 0, 1 and 2 compared to all upregulated cluster markers.
E. Module score depicting the average expression of the significantly upregulated genes from the RNA sequencing analysis (fig. 3F) in the individual stromal cell clusters.
F. GO Enrichment analysis for upregulated genes in stromal cell clusters 0, 2 and 5 compared to all upregulated cluster markers.

**Supplementary Figure 7**

A. Analysis of the Fold Enrichment of the GO term lipid metabolic process (GO:0006629), including its child GO terms in the stromal cell clusters.
B. Analysis of the Fold Enrichment of the GO term autophagocytosis (GO:0006914), including its child GO terms in the stromal cell clusters.
C. C. Heatmaps of CellChat analysis depicting sender-recipient interactions between macrophage and stromal cell subsets in control and infection.
D. D. Bubble plots depict the probability of ligand-receptor signaling interactions between stromal and macrophage clusters for the top nine predicted pathways (probability ≥0.05) in control and infection.
E. Supplementary Figure 8
F. A. Depicted are means ± SEM of absolute numbers of macrophage subsets per omentum in mice from 6B (corresponds to Figures 6D-G)
G. B. Depicted are means ± SEM of percentages of macrophage subsets in omentum in mice from 6B.
H. C. Shown are percentages of iNOS^+^ cells of F4/80^+^ CD11b^+^ cells in omentum in mice from 6B.
I. D. Depicted are means ± SEM of spleen and body weights (compare 6C).
J. E. Depicted are means ± SEM of SPM and Ly6C^+^ monocytes in peritoneal lavage of the mice in 6C.
K. CFU counts in spleen, liver and BM from mice in 6C.
L. G. NicheNet analysis from fig. 6M comparing the ligand expression on stromal cells to differentially expressed genes (receptors) between inflammatory monocytes of high and low burden granulomas.

## METHODS

### Mice

All mice were on C57BL/6 genetic background. We included both male and female mice, usually at the age of 6 to 12 weeks, in the experiments. C57BL/6J mice were purchased from the CEMT animal facilities (University of Freiburg). *Pdgfrα^CreERT^*^2^ mice (B6.129S-Pdgfratm1.1(cre/ERT2)Blh/J) and β-Actin-GFP mice (C57BL/6-Tg(CAG-EGFP)131Osb/LeySopJ) were purchased from Jackson Laboratories (USA). Rebecca Kesselring (Department of General and Visceral Surgery, Medical Center, University of Freiburg) provided β-Actin-dsred mice (B6.Cg-Tg(CAG-DsRed*MST)1Nagy/J) as a kind gift. β-actin mice were used both homozygous and heterozygous for the experiments as we did not observe relevant differences in fluorochrome expression. *Cxcr4^CreER^*mice were provided by Ralf Stumm (Neuropharmacology, Jena) and LifeAct-GFP mice by Tim Lämmermann (Zentrum für Molekularbiologie der Entzündung, University Münster) as kind gifts. Mice for infection experiments at biosafety level 2 were bred in the CEMT animal facilities of the University of Freiburg in groups of maximum 5 animals under specific pathogen-free conditions. The room temperature was set to 21°C (± 0.5-1°C), the humidity to 55 % (± 5-8 %) and day/night cycles to 12 hours. Food and water were provided ad libitum. LifeAct-GFP mice were kept at the Max Planck Institute of Immunobiology and Epigenetics (Freiburg). Animal experiments under Biosafety level 2 were approved by the Regierungspräsidium Freiburg, Germany (G-19/171, G-22/097).

With the aim of minimizing the number of experimental mice required for *M.tb* infections, animals from another simultaneously conducted *M.tb* infection experiment were used to extract pleural lavage fluid. For this reason, mice of the strain Ebi3^Tom.L/L.Thy1.1^ (B6-Ebi3tm2(Tom/Thy1.1)Vig/J)^85^ were used, which are on a C57BL/6 genetic background and behave phenotypically equivalent to wild-type mice. Breeding of those animals took place at the Saxon Incubator for Clinical Translation (Leipzig, Germany). The uninfected control group additionally includes C57BL/6N mice of a similar age, which were purchased from Charles River. All mice were kept under barrier conditions in individually ventilated cages. The experiment was performed in the BSL 3 facility at the Research Center Borstel (Borstel, Germany) according to the German animal protection laws and was approved by the Animal Research Ethics Board of the Ministry of Agriculture, Rural Areas, European Affairs and Consumer Protection (Kiel, Germany) (approval number 19-3/21).

### Tamoxifen induction

*Cxcr4^creER^:R26^tdTomato^* received tamoxifen subcutaneously twice 4 weeks prior to i.p. infection. Tamoxifen (Sigma Aldrich) was dissolved in Corn oil (Sigma Aldrich) at 4 mg/200 µl by gentle heating (37-42°C) and shaking over several hours. Subcutaneous injections (50 µl per limb; total 4 mg per administration) were performed under isoflurane anesthesia on all four limbs, twice, with a 2-day interval. *Pdgfrα^CreERT2^:R26^DTA+/-^*or *R26^DTA-/-^* received tamoxifen orally by micropipetting. A 100 mg/ml tamoxifen stock in peanut oil (Sigma-Aldrich) was prepared by dissolving tamoxifen initially in 100% ethanol, overlaying with peanut oil, then fully evaporating ethanol at 55 °C with shaking. The stock was emulsified 1:1 with a sweetened condensed milk–water mixture (1:2) using a Luer–Luer-lock system, as previously described^86^. Starting one day before i.p. infection (d-1), mice were trained for voluntary oral uptake of the oil–milk emulsion without tamoxifen over 3 consecutive days. On day 2 p.i., mice received tamoxifen–milk emulsion with 3 mg tamoxifen daily for 5 days. While training and the first tamoxifen dose were voluntarily consumed, mice were restrained for subsequent administrations.

### Preparation of *M. bovis* BCG

*M. bovis* BCG was provided by Prof. Dr. Dirk Wagner (Clinic for Internal Medicine II, Gastroenterology, Hepatology, Endocrinology and Infectious Disease, Medical Center - University of Freiburg) as a kind gift. To generate BCG-BFP, 100 µl competent BCG was electroporated (2500 V, 25 µF, 1000 Ohm) with 1 µg DNA of a purified BFP plasmid (Addgene plasmid # 30177; http://n2t.net/addgene:30177; RRID:Addgene_30177)^87^. BCG was grown in liquid culture of 7H9 broth with 10 % OADC and 0.5 % glycerol. After 24 h, 50 µg/ml hygromycin was added for selection and cultures were grown to an OD600 of 1. Afterwards, the liquid culture was pelleted and stored in aliquots of 7H9 broth with 10 % OADC and 15 % glycerol at -80 °C till usage.

### Intraperitoneal infection

*M. bovis* BCG was resuspended in PBS and homogenized with a 28G syringe and by sonication. The mice were infected i.p. with ∼1.6-1.8 * 10^7^ CFU in 100 µl PBS. PBS alone served as vector control. The injection was preferably administered in the right lower abdominal quadrant.

### Tissue preparation and digestion

Mice were sacrificed in CO_2_ and blood was collected immediately after death from the retroorbital plexus in heparine and serum tubes. To analyze the blood cells in flow cytometry an RBC lysis (eBioscience™, 1x RBC lysis buffer) was performed twice 5 minutes at room temperature. After centrifugation (3600 rpm, 6 min) the supernatant was removed carefully and the pellet then washed in FACS buffer. Alternatively, a Fixative-Free lysing solution (High-yield lysis buffer) was used after FACS staining for 10 minutes at room temperature. After washing, the cells were resuspended in the desired amount of FACS buffer. To gain the serum, the serum tubes were kept for at least 30 minutes at room temperature that the blood can coagulate. The samples were then centrifuged at 2000 g for 10 minutes and the serum carefully removed and stored at -20°C for further processing.

For the peritoneal lavage the skin of the abdomen was carefully incised without damaging the peritoneal membrane. 5 ml FACS buffer were injected carefully with a 27G needle into the peritoneal cavity and the liquid removed after a small incision with a 1 ml pipette. The pleural lavage was performed by injecting small amounts of buffer (approximately 0.5 ml -1 ml) via the diaphragma and again removed after a small incision with a 1 ml pipette. This process was repeated several times. The lavages were centrifuged and the cell pellets resuspended in the desired amount of FACS buffer for further processing.

The omentum is localized below the stomach and was carefully removed. If needed, parts of the omentum were saved for microscopy. The remaining parts were digested in 5 ml RPMI with 2% FBS supplemented with collagenase 2 (Worthington, 1 mg/ml), DNase 1 (Sigma-Aldrich, 0.5 mg/ml) and dispase (Corning, 354235, 0.05 ml/ml) for 30 minutes in a water bath at 37°C. The tissue lysate was vigorously vortexed several times during and after enzymatic digestion and then mechanically smashed through a 70 µm cell strainer to obtain a single-cell suspension. Dense cores of highly mature granulomas, which could not be fully dissociated by digestion and mechanical disruption, were retained on the filter and excluded from downstream analysis. The resulting cell suspension was centrifuged at 1600 rpm for 6 minutes at 4°C. The supernatant was discarded, and the cell pellet was washed once with FACS buffer (PBS, 1% BSA, 2 mM EDTA) before proceeding to analysis. After a second centrifugation the pellet was resuspended in the desired amount of FACS buffer. For determination of absolute cell counts, the omentum was weighed prior to enzymatic digestion, then fully digested and resuspended in a defined volume of 200 µl FACS buffer.

### Oropharyngeal infection and transfer

For the oropharyngeal infection mice were shortly put under isoflurane narcosis and up to 5 * 10^7^ BCG in 45 µl were installed into the throat with a pipette while the tongue was slightly fixed. Control mice received PBS alone. 2 days later stromal cells were isolated from the omentum of β-Actin-GFP donor mice as described above and approximately 1,2-1,5 * 10^5^ SC were applied oropharyngeally in 45 µl PBS as described. At the analysis time point mice were sacrificed in CO_2_ and perfused with PBS post mortem, when FACS analysis of the lung was performed. The lung was inflated with 700 µl of dispase (Corning, 354235) and cooled in PBS till further processing. Digestion buffer (0,5% BSA in PBS) with collagenase 1 (Worthington, 5 mg/ml) and DNase (Sigma-Aldrich, 1 mg/ml) was prepared freshly. Lungs were cutted into 1-2 mm pieces in 2 ml digestion buffer and incubated on a horizontal shaker 30 min at 37°C, 130 RPM. After digestion, the suspension was homogenized with a syringe (18G needle, 5 ml syringe) and smashed through a 70 µm strainer. After washing, and a 5 min RBC lysis, each lung cell suspension was resuspended in 1 ml FACS buffer for further processing. If lungs were used for immunofluorescence staining, mice were perfused with 4% PFA post mortem and lungs were slowly inflated with 700 µl PFA to preserve the lung architecture. Lungs were kept in PFA overnight till further processing.

### Histology

Tissues were fixed in 4% PFA for 24 hours and H&E stainings were performed with the Tissue-Tek Ready-to-use kit (Sakura, Hematoxylin 9130-E, Eosin 9134-E). In short, samples were dried and deparaffinized with Xylol twice 3 minutes. After performing a descending ethanol series, slides were washed and stained in Hematoxylin for 5 minutes. After 3 minutes of washing samples were put into 70% ethanol for 30 seconds and stained with eosin for 3 minutes. An ascending ethanol series was performed and samples incubated in Xylol for 2 and 3 minutes. PAS (Periodic Acid-Schiff), AFOG (Acid Fuchsin-Orange G) and EvG (Elastica-van-Gieson) stainings were performed on a Prisma staining device according to manufactureŕs instructions (Sakura, Finetek, Europe). For semiquantitative scoring of lymphofollicular structures respective samples were assessed (20x magnification level) and scored (following criteria were defined: 0 – no (organized) lymphoid follicles apparent, 1 – minimal, residual lymphoid follicles discernible, 2 – normal, well-developed lymphoid structures; at least n=3 animals per time point were assessed). The sections were scanned with a Ventana 200 D scanner (20x).

### Paraffin blocks and immunohistochemistry staining of paraffin slides

Lungs were kept in 4% PFA for at least 24 hours and afterwards paraffin embedded with the Sakura Tissue-Tek VIP 6 AI. 4 µm cuts were performed for immunofluorescence staining. The samples were incubated in citrate buffer for 5 minutes and afterwards blocked in 5% goat serum/ 5% bovine serum for 30 minutes. Primary antibody staining with rat Nos2 PE 1:50 (eBioscience), rabbit F4/80 1:200 (D2S9R, cell signaling) and goat GFP DyLight 1:1000 (Thermo Fisher Scientific) was performed overnight. After several washing steps with 1x Washing buffer (S3006, DAKO), secondary staining with anti-rabbit Alexa Fluor 647 1:500 (4 µg/ml) and nuclear staining with Hoechst 33342 1:1000 (250 µg/ml) was performed for 45 minutes. For antibody dilution an antibody diluent was used (S3022 + 2.5% BSA, DAKO). After secondary staining samples were washed and mounted with ProLong mounting medium Invitrogen).

### Isolation of monocyte progenitors from bone marrow

Bone marrow progenitors were isolated as previously described^16^. In short, bones were flushed with sort buffer (PBS with 1% BSA, 2 mM EDTA) through a 70 µm strainer and stained with biotinylated Ter119, CD3e, CD19, SiglecF, CD127, Ly6G antibodies at 4°C for 20 minutes. Biotin-labelled cells were removed with Dynabeads™ (Thermo Fisher Scientific). Progenitor cells were stained with CD115, CD117, CD45 (except for β-Actin-GFP mice), CD135, Ly6C and CD11b. iMoP were sorted as CD115^+^, CD117^-^, CD45^+^, CD135^-^, Ly6C^+^, CD11b^-^. Bone marrow progenitors were isolated on a MoFlo® Astrios™ cell sorter (Beckman Coulter).

### Adoptive transfer of LPM i.p

Peritoneal lavage from β-Actin-GFP mice was stained for F4/80 PE, CD11b Pe-Cy7, CD115 BV421, Ly6C BV605, MHC-II APCeFl 780 and LPM (F4/80^+^, CD11b^+^, CD115^+^, MHC-II^-^, Ly6C^-^) were sorted into FACS buffer (PBS + 1% BSA + 2 mM EDTA). Cells were centrifuged (1600 RPM, 6 min, 4°C) and resuspended in PBS. 4.5 * 10^5^ LPM in 100 µl PBS were injected i.p.. Infection i.p. (respectively PBS injection) was performed at the same time point. Mice were sacrificed 24 hours post transfer and omentum and peritoneal lavage analyzed for transferred LPM.

### Adoptive transfer of monocyte progenitors i.p

Recipient mice were i.p. infected 14 days prior to adoptive transfer. For isolation of monocyte progenitors from β-Actin-GFP mice, we followed the above isolation protocol. For the final Sort staining we used CD115 REAlease® PE antibody, CD11b APCeFl780, Ly6C BV605 and sorted for CD115^+^, CD11b^-^, Ly6C^+^ and GFP^+^ cells, which includes cMoP and iMoP. The REAlease® kit (Miltenyi Biotec) was used to remove CD115 antibody according to the manufactureŕs instructions. 3-4 * 10^5^ monocyte progenitors were injected i.p. in 100 µl PBS and mice were analyzed 2 weeks post adoptive transfer (4 weeks p.i.).

### Adoptive transfer of stromal cells i.p

Omentum from β-Actin-GFP mice was digested as described above and stained for CD45 PE; CD31 PE, PDPN AF647. Cells were isolated on a Cytoflex SRT (Beckman Coulter). 0.7-2.5 * 10^5^ GFP^+^, PDPN^+^, CD31^-^ SC were injected into recipients and analysis was performed on day 1 or 3 post transfer. At the 3 days analysis time point omentum was digested and described and GFP^+^ transferred SC were sorted from the recipients. The sorted cells were cultured in µ-Slide 8 Well high (ibidi, 80806) and microscopically analyzed for intracellular BCG-BFP. The number of bacterial positive cells was set into relation to the number of plated cells.

### Antibiotic treatment

*Cxcr4^creER^:R26^tdTomato^* mice were induced with tamoxifen subcutaneously as described above. 4 weeks later BCG/PBS i.p. injections were performed. 2 weeks p.i. oral antibiotic treatment was started for another 4 weeks. Rifampicin and isoniazid (Sigma-Aldrich) were both diluted at 100 mg/l in the drinking water. Antibiotics were refreshed every 2 days and the drinking bottles weighed to ensure intake.

### Omentum whole mount

For microscopy of omentum whole mounts, small pieces of the omentum were incubated after preparation for 1 hour in 4% PFA. The tissue was then transferred into blocking buffer (PBS, 1% BSA, 0.1% Triton-X) for at least 30 minutes. The tissue was stained in blocking buffer over night with the respective antibodies. Afterwards samples were washed in blocking buffer three times for 5 minutes. After washing, the mounting followed in Prolong Diamond Mounting Medium (∼2 drops). Whole mounts were tried for at least 24 hours before confocal imaging was performed on LSM 880 confocal microscope with a ×20 objective (numerical aperture 0.8) or a x40 objective (numerical aperture 1.2) and analysis followed with ZEN.

### Granuloma imaging

For Granuloma immunofluorescence staining, granuloma structures were isolated from omentum and put into 4% PFA for 4 hours. Tissue samples were transferred into 20% sucrose till they sank. Granulomas were cryo-embedded in Tissue-Tek® O.C.T™ Compound (Sakura) and shock frozen in liquid nitrogen. Slices of 8 to 10-μm thickness were cut at a cryotome. Slides were washed with washing buffer (0.1% Triton-X100, 1% BSA in 1× PBS) and kept in the buffer for blocking for 5-6 hours at 4°C in the dark. Slices were stained with Phospho-Histone H2A.X (Ser139) (20E3) Rabbit mAb #9718 1:200 (Cell Signaling) and F4/80 PE (Invitrogen, 1:50) overnight in the dark at 4°C. Slides were washed 3 times in washing buffer. Samples were stained with Acti-stain™ 488 Fluorescent Phalloidin (Cytoskeleton, 1:250) and anti-rabbit secondary antibody 633 1:500 for 1 hour at room temperature. For the neutrophil staining (Fig. 1F), overnight incubation was performed with or Ly-6G PE (BD Pharmigen), F4/80 AF488 (BioLegend, 1:100) and Podoplanin (BioLegend, 1:100). Granuloma sections were washed 3 times and mounted with ProLong™ Diamond Antifade Mountant (invitrogen). Samples were tried for 24 h at room temperature before imaging or storage at 4°C. Tile scans were acquired on LSM 880 with a ×20 objective (numerical aperture 0.8) and an overlay of 5% (Fig. 1) or 10% (Fig. 1F). Images were stitched in ZEN. Maximal intensity projection was performed for the stainings in (Fig. 1F).

### CFU counts

To evaluate CFU counts bone marrow of one leg (femur, hip, tibia) was flushed through a 70 µm filter and resuspended in 300 µl PBS. A piece of liver tissue and spleen were weighed and passed through a 70 µm strainer in 1 ml PBS each. The omentum (Supplementary Fig. 1B) was homogenized twice for 5 minutes each in 500 µl of PBS at 50 oscillations per second using a homogenizer. Serial dilutions were performed on Middlebrook 7H10 agar plates, growth was regularly evaluated and colonies counted for 4-6 weeks.

### FACS staining

For FACS analysis samples were blocked with anti-FcR antibody (1:100) for 5-10 minutes on ice. FACS analysis was in general performed in 100 µl FACS buffer. The antibody concentrations were adjusted individually. Antibody stainings were performed for 30 minutes at 4°C and samples then washed by addition of at least 1 ml FACS buffer. After centrifugation the supernatant was discarded and the pellet resuspended in 200 µl FACS buffer for acquisition at a 3-laser flow cytometer (Gallios™, Beckman Coulter). FACS data were processed with the Kaluza software (v2.3, Beckman Coulter). If the primary antibody is biotin-conjugated a second staining step with a streptavidin antibody followed.

For intracellular staining (Nos2, TNF) first the surface staining was performed as described. After washing samples were fixed and permeabilized with the Fixation/Permeabilization Solution Kit (BD Bioscience) according to manufacturer’s instructions. Pleura fluid from *M.tb* infected mice was acquired on a MACSQuant Analyzer 10 (Miltenyi Biotec).

### Antibodies

The following mouse antibodies were used: PDL1 APC (10F.9G2, #124312), Ly6C BV605 (HK1.4, #128035), CD115 BV421 (AFS98, #135513), Podoplanin AF647 (PMab1, # 156204), CD3e AF488 (17A2, #100212), Ly6G APC (1A8, # 127614), F4/80 AF488 (BM8, # 123120), CD31 PE (390, #102407), CD31 FITC (390, #102405), PDGFRA Biotin (APA5, #135909), Ly6G Biotin (1A8, #127604), CD117 BV421 (2B8; #105828), CD68 AF647, (FA-11; #137004), CD45 FITC (30-F11, #103108), CD45 BV605 (30-F11, #103139), SiglecF FITC (S17007L, # 155503), SiglecF APC (S17007L, #155508), CD135 (A2F10, #135309), CD64 PerCP-Cy5.5 (X54-5/7.1, #139308), Ep-CAM PerCP-Cy5.5 (G8.8, #118219), CD16/32 purified (93; #101301), Precision Counting Beads (#424902) were purchased from BioLegend.

CD11b PE-Cy7 (M1/70, #25-0112-82), CD11b APC-eFluor 780 (M1/70; #47-0112-80), CD45 FITC (30-F11, #11-0451-82), CD45 PerCPcy5.5 (30-F11, #45-0451-82), CD45 eFluor 450 (30-F11; #48-0451-82), F4/80 PE (BM-8, #12-4801-82), TNF PE (MP6-XT22, #12-7321-82), TIM-4 AF488 (RMT4-54, #53-5866-80), TIM-4 AF647 (RMT4-54, #130008), NOS2 AF488 (CXNFT, #53-5920-80), NOS2 APC (CXNFT, #17-5920-80), NOS2 PE (CXNFT, #12-5920-80), Ly6G PE (1A8, #127607), CD3e PE (145-2C11, #12-0031-82), MHC-II APCeFluor-780 (M5/114.15.2, #47-5321-82), CD31 PE (390, #12-0311-81), Ly6C PerCP-Cy5.5 (HK1.4, #45-5932-82), Streptavidin PE-Cy7 (#25431782), anti-GFP DyLight 488 (Goat polyclonal; #600-141-215), Alexa Fluor 633 goat anti rabbit (#A21071), Alexa 647 goat anti rb (#A21245) were obtained from Thermofisher Scientific.

CD115 PE (AFS98, #130-122-018), CD31 FITC (390, #1301-236-75), Ter119 Biotin (Ter119, #130-120-828), CD3e Biotin (145-2C11, #130-101-990), CD19 Biotin (6D5, #130-119-797), SiglecF Biotin (ES22-10D8, #130-119-136), Sca-1 Biotin (D7, #130-126-949), CD127 Biotin (A7R34 and REA680, #130-102-039 and #130-110-373), CD115 PE REAlease (REAL272, #130-118-071) and CD115 Antibody Kit, PE, REAlease® (#130-122-632) were bought from Miltenyi Biotec and CD11c FITC (HL3, #553801), TNF APC (MP6-XT22; #554420), Ly6G PE (1A8, #551461), CD103 PE (M290, #561043) from BD bioscience. Phospho-Histone H2A.X (Ser139) (20E3) (Rabbit mAb #9718) and F4/80 (D2S9R rb MAB #70076S) were purchased from Cell Signaling. Ly-6G PE antibody was bought from BD Pharmigen (1A8, # 551461).

### Aldefluor assay

Aldefluor assay was performed according to manufactureŕs instructions (AldeRed ALDH Detection Assay, Merck). In short, digested omentum samples at 24 hours and 4 weeks p.i. as well as controls were resuspended in 500 µl assay buffer supplemented with verapamil. 2.5 µl AldeRed 588 was added to the suspension and 250 µl of the solution directly transferred to a control tube with 2.5 µl DEAB (background control). The suspensions were incubated 60 min at 37°C and centrifuged at 250 g for 5 minutes. Samples were stained in 100 µl assay buffer with Podoplanin AF647 and CD45 FITC for 30 minutes on ice. After washing with 500 µl assay buffer, cells were resuspended in 150 µl assay buffer and acquired on a 3-laser flow cytometer (Gallios™, Beckman Coulter).

### Ziehl-Neelsen-Staining

Granuloma cryosections were washed in PBS and heat-fixed for 10 min at 100°C. The slide was covered with carbolfuchsin staining and incubated on a heating coil for 15 minutes. Samples are washed with H_2_O and decolorized with HCl-alcohol (approx. 2 minutes). After washing, samples are restained with methylenblue for 20 seconds. Samples were washed with H_2_O and tried.

### RNA extraction and quantitative real-time PCR

Cell lysis was performed in RLT lysis buffer (Qiagen) with 1% β-mercaptoethanol and RNA extracted with RNeasy MicroKit (Qiagen) according to the manufacturer’s instructions. Superscript IV (Invitrogen) was used to transcribe RNA into cDNA. RT-qPCR was conducted with absolute qPCR SYBR Green Mix (Thermo Fisher Scientific) in a LightCycler 480 systems (Roche). mRNA levels were normalized to the arithmetic mean of ct values of *Gapdh*, *Hprt* and *Actin* as housekeeping genes.

### Primer Sequences

The following mouse primers were used for RT-qPCR (5‘-3‘):

- *Actin* (for: AGATCAAGATCATTGCTCCTCCT, rev: ACGCAGCTCAGTAACAGTCC)
- *Gapdh* (for: ACTCCACTCACGGCAAATTC, rev: TCTCCATGGTGGTGAAGACA)
- *Hprt* (for: CTTTGCTGACCTGCTGGATT, rev: TATGTCCCCCGTTGACTGAT)
- *Raldh1* (for: ATGGTTTAGCAGCAGGACTCTTC, rev: CCAGACATCTTGAATCCACCGAA)
- *Raldh2* (for: GACTTGTAGCAGCTGTCTTCACT, rev: TCACCCATTTCTCTCCCATTTCC)
- *Tlr2* (for: CACCACTGCCCGTAGATGAAG, rev: AGGGTACAGTCGTCGAACTCT)
- *Ccl2* (for: AGCCAACTCTCACTGAAGCC, rev: GCTTGGTGACAAAAACTACAGC)
- *Ccl7* (for: CCTCTTGGGGATCTTTTGTTC, rev: ATCTCTGCCACGCTCTGTG)
- *Abca1* (for: TGG AAA CTC ACC CAG CAA CA, rev: GGC AGG ACA ATC TGA GCA AAG)
- *Dhcr24* (for: GAGGGCTTGGGATACTGCAC, rev: CCTTCACAGGCCAAATGGATG)
- *Fasn* (for: GGCTCTATGGATTACCCAAGC, rev: CCAGTGTTCGTTCCTCGGA)

### Smart-3 Sequencing

SC were sorted as described above and cultured in full Opti-MEM with 10% FBS supplemented with 1% Ciprofloxacin. SC were infected with BCG MOI12. Uninfected SC served as control. Cells were lyzed at day 3 of culture in RLT buffer and RNA purified with the RNeasy MicroKit (Qiagen). RNA sequencing libraries were prepared using the Smart-3SEQ protocol with minor modifications^88^. Briefly, total RNA was fragmented in SMARTScribe first-strand reaction buffer with (Takara Bio) dNTPs (New England Biolabs), followed by reverse transcription using an anchored oligo(dT) primer. Template switching was performed using SMARTScribe reverse transcriptase (Takara Bio) to add a 5’ adapter sequence directly to the cDNA. The resulting cDNA libraries were amplified by PCR using indexed primers to complete the Illumina-compatible adapters (Takara Bio). Amplified libraries were purified using SPRI beads (Beckman Coulter Life Sciences) to remove adapter dimers and size-selected for cDNA inserts. Final libraries were quantified and pooled for sequencing on an Illumina NextSeq 1000/2000 platform. Sequencing reads were aligned to the Gencode vM33 reference genome, and gene expression was quantified using a custom script in usegalaxy.eu. A detailed experimental protocol can be found here: <u>DOI:</u> dx.doi.org/10.17504/protocols.io.81wgbwx2ygpk/v1

The downstream analysis was performed in R (version 4.5.1). The DESeq analysis was performed using the DESeqDataSetFromMatrix function using the DESeq2 package^89^. To convert ENSEMBL IDs, we used the gencode.vM33.annotation.gtf.gz (https://www.gencodegenes.org/mouse/releases.html) with the BioMatr package^90^ and org.Mm.eg.db package. Prior to differential expression analysis, row sum counts >=10 were selected. Size factor normalization was performed using DESeq2’s estimateSizeFactors function to adjust for differences in sequencing depth across samples. Normalized counts were then extracted from the DESeqDataSet. Variance stabilizing transformation (VST) was applied to the normalized counts using vst with blind = FALSE to generate log2-like transformed data suitable for downstream analysis. PCA plot was generated using the plotPCA function. Differential expression was performed with the DESeq() and results for infected versus control SC were extracted using the results function. Log2 fold changes were then shrunk via the apeglm method to improve effect size estimation. We selected the relevant differential expressed genes (baseMean > 10, log2FC > 1 and padj < 0.05) for plotting and analysis. The GO term analysis was performed with the PANTHER^91^ Overrepresentation Test (Released 20231017), GO Ontology database DOI: 10.5281/zenodo.10536401 Released 2024-01-17, FDR correction. The results (dds) gene list was used as background gene list.

### Single Cell Sequencing of omentum macrophages and stromal cells

Single-cell RNA sequencing was performed using 10x Genomics with feature barcoding technology to multiplex cell samples from different conditions to reduce costs and minimize technical variability. Mice were infected i.p. as described 4 weeks prior to the sequencing. We used 3 BCG-infected and 5 PBS control mice for the analysis. FACS staining was performed for PDPN, CD11b, CD45, F4/80, CD31, Ly6G and Zombie Green. The individual samples were marked by 10x Genomics 3’ CellPlex Multiplexing Solution as followed: PBS1 CM301, PBS2 CM310, PBS3 CM303, PBS4 CM304, PBS5 CM305, BCG1 CM306, BCG2 CM308, BCG3 CM309. Each control and infected samples were pooled. Mφ were sorted as Lin^-^/Live (CD31, Ly6G, Zombie), CD45^+^, CD11b^+^, F4/80^+^ and SC as Lin^-^/Live, CD45^-^, PDPN^+^. Sorted cells were processed through the 10x Genomics 3′ v3.1 workflow according to the manufacturer’s instructions. Libraries were pooled to desired quantities to obtain appropriate sequencing depths as recommended by 10x Genomics and sequenced on a NovaSeq 6000 flow cell. Quantification of gene expression and CellPlex oligonucleotide tags was performed using cellranger-8.0.0 using the count command, which performs alignment, filtering, barcode counting and UMI counting as well as process feature barcoding data. Alignments were performed using prebuilt Cell Ranger mouse reference (refdata-gex-GRCm39-2024-A).

The downstream analysis was performed in R (version 4.5.1). We used the Seurat package (version 5.3.1^92^) and for demultiplexing HTODemux with a positive quantile threshold of 0.99 was performed. The analysis was proceeded with 19,813 singlets for Mφ and 19,183 singlets for SC. For the Mφ SCS, cells with total RNA counts (nCount_RNA) greater than 15,000 or with more than 10% mitochondrial gene content (percent.mt) were excluded. The remaining cells (control: 7,062; infected: 9,240) were further filtered to retain those with more than 500 detected genes (nFeature_RNA). For the stromal cell SCS, cells with total RNA counts (nCount_RNA) greater than 12,000 or with more than 10% mitochondrial gene content (percent.mt) were excluded. The remaining cells (control: 10,098; infected: 7,134) were further selected to retain those with more than 500 detected genes (nFeature_RNA). As we aimed for a high Mφ purity for our further downstream analyses, we removed cells with a gene count threshold > 1 for the contaminating genes (*Cd19, Ncr1, Pdgfra, Cd3d, Cd3g, Cd3e, Cd79a, Zbtb46, Myom3, Ptprcap, Cpa1, Ebf1, Cpa3, Pou2af1, Wt1, Skap1, Upk3b*). We normalized with the NormalizeData() function and the data was further subjected to FindVariableFeatures(), ScaleData(), and RunPCA() algorithms. FindNeighbours and FindClusters functions were applied with the top 30 harmony dimensions and a resolution of 0.5. We used the FindAllMarkers() function with a min.pct of 0.25, a logfc.threshold of 0.25 and an adjusted p-value of 0.05. Th e contaminating and proliferating clusters 4, 7, 8, 9 and 10 (initial cluster numbers) were removed. The normalization and reclustering with a resolution of 0.5 was reperformed as described.

For the stromal cell SCS, we removed cells with a gene count threshold > 1 for the contaminating genes (*Ptprc, Epcam, Cd209a, Cd209b, Cd209f, Cd33, Amy2a3, Adgre1, Cpa1*). Initial normalization and clustering was performed as described for Mφ with a resolution of 0.3 for FindClusters(). The proliferating and contaminating clusters 9 and 10 (initial cluster numbers) were removed. The normalization and reclustering was repeated with a resolution of 0.25.

The clusters were depicted as DimPlots. The cluster annotations were performed based on the differentially upregulated genes with the FindAllMarkers() function (min.pct 0.25, logfc.threshold 0.25, adjusted p-value of 0.05). Heatmaps were depicted with the DoHeatmap() function of the pheatmap package (version 1.0.13^93^) after scaling the datasets and dotplots of selected genes with the DotPlot() function. To compare the results of the Smart-3 Sequencing analysis with the single-cell sequencing dataset, module scores were calculated for each stromal cell cluster using Seurat’s AddModuleScore function, based on genes significantly upregulated in infected SC. The distribution of module scores across clusters was visualized using violin plots generated with ggplot2^94^ (version 4.0.0). We calculated the enriched GO terms for selected clusters with the enrichGO function of the clusterProfiler^95^ package (version 4.18.1) with ont = BP (biological process), pAdjustMethod = BH and a qvalueCutoff = 0.05. In addition, we calculated if specific GO terms (autophagocytosis GO:0006914; endocytosis GO:0006897; lipid metabolic process GO:0006629) and their child terms were significantly enriched in the SC clusters, visualizing fold enrichment values per cluster as heatmaps. For the CellChat interaction analysis, we used the CellChat package^96^ (version 2.2.0), performing separate analyses for control and infected Mφ-stromal cell interactions. Clusters with fewer than 30 cells were removed from the Seurat objects. The control Mφ and stromal Seurat objects and, separately, the infected Mφ and stromal Seurat objects were each merged. The CellChat workflow—including createCellChat, identifyOverExpressedGenes(), identifyOverExpressedInteractions(), subsetData(), computeCommunProb(), computeCommunProbPathway(), and aggregateNet()—was then performed. Results were visualized as heatmaps using netVisual_heatmap(). Bubble plots were generated for the nine most relevant pathways with a communication probability ≥ 0.05, considering SC as senders and Mφ as targets. Finally, interaction networks between SC (senders) and Mφ (targets) for CCL signaling were depicted using the netVisual_chord_gene() function. For the NicheNet analysis we used the nichenetr package^97^ (version 2.2.1.1). First, we calculated the significantly upregulated genes between infected and control Mφ with the FindMarkers function and a min.pct of 0.1, filtered for and adjusted p-value of <0.05 and a log2FC of >0.5 (padjust < 0.05, log2FC > 0.5), which we defined as target genes. Ligand–receptor network data were obtained from the Zenodo repository associated with NicheNet (accessed at https://zenodo.org/record/7074291). Stromal and Mφ cell expression profiles were used to define expressed ligands and receptors (expressed in ≥10% of cells). Ligand activities were inferred using the predict_ligand_activities function, and top ligands were visualized with ggplot2. For all sequencing analysis, the dplyr package (version 1.1.4)^98^ was used.

### Single cell sequencing reanalysis of the Gideon et al.^9^ dataset

We used the GSE200151 data set of *M.tb* infected lungs of Macaca fascicularis 10 weeks p.i. together with the published metadata. We assigned the metadata and performed the Seurat workflow with standard settings as described above. For the NicheNet analysis we calculated the upregulated genes of the cluster annotated as Mφ (Mphage) between high and low burden granulomas with a min.pct of 0.1., filtered for and adjusted p-value of <0.05 and a log2FC of >0.5, which we defined as target genes. Ligand–receptor network data were obtained from the Zenodo repository associated with NicheNet for human (accessed at https://zenodo.org/record/7074291). Fibroblast and macrophage cell expression profiles were used to define expressed ligands and receptors (expressed in ≥10% of cells). Ligand activities were inferred using the predict_ligand_activities function, and top ligands were visualized with ggplot2.

### Single cell sequencing reanalysis of the Wang et al.^45^ dataset

We used the GSE192483 human data sets from tuberculosis patients with low and high burden. We used the CreateSeuratObject function with min.cells =3, min.features > 200 and < 5000. First the standard Seurat workflow was performed on the low-burden and high-burden data set separately with standard settings. The data sets were merged and filtered for a percent.mt < 15. The standard Seurat workflow was repeated on the merged object with a resolution of 0.3. Marker genes for each cluster were identified using Seurat’s FindAllMarkers function with parameters only.pos = TRUE, min.pct = 0.25, and logfc.threshold = 1. Markers were then filtered to retain those with average log2 fold change > 1. A composite score (score = avg_log2FC * log10(1 / (p_val + 1e-300)) * pct.1) was calculated to rank markers. The top 50 markers per cluster were selected and the annotation of clusters of interest (Mφ, inflammatory monocytes and SC) based on that analysis. For the NicheNet analysis we calculated the upregulated genes of the Mφ clusters and inflammatory monocyte cluster between high and low burden each with a min.pct of 0.1., filtered for and adjusted p-value of <0.05 and a log2FC of >1, which we defined as target genes for the three individual NicheNet analyses. Ligand–receptor network data were obtained from the Zenodo repository associated with NicheNet for human (accessed at https://zenodo.org/record/7074291). Fibroblast and Mφ cell expression profiles were used to define expressed ligands and receptors (expressed in ≥10% of cells). Ligand activities were inferred using the predict_ligand_activities function, and top ligands were visualized with ggplot2.

### *M.tb* infection

For conducting the infection experiment, a stock solution of *M.tb* H37Rv was diluted in sterile distilled water and pulmonary infection of experimental mice was performed using an inhalation exposure system (Glas-Col, Terre-Haute, IN, USA). Thereby, mice were exposed for 40 min to an aerosol generated by nebulizing 5.5 ml of a suspension containing approximately 10^7^ live bacteria. Twenty-four hours post-infection, the inoculum size was controlled by determining the bacterial load in the entire lung of infected mice. Animals were infected with an actual infectious dose of 122 CFU/lung. The mice were infected for 42 and 84 days before the pleura lavage was performed, as described above.

### Bacteria and particle uptake

SC were isolated from WT mice as PDPN AF647^+^ CD31 PE^-^ CD45 FITC^-^ and 20,000 SC plated in 12 well plates in 1 ml Opti-MEM supplemented with 10% FBS either with or without ciprofloxacin and BCG-BFP (MOI12) added. Uninfected cells served as control. Additionally, separate conditions were treated with biotin-coated yellow fluorescent particles (2.2 µl, 0.47 µm, Spherotech). After 3 days of culture, the supernatant was removed and cells were stained with PDPN AF647 and Streptavidin PE-Cy7. The uptake of particles was evaluated in FACS as on the one hand BFP-positive SC and on the other yellow fluorescent, PE-Cy7^-^ SC.

### Bacterial survival assay

SC were sorted from WT Omentum as PDPN^+^ CD31^-^ CD45^-^ cells and 5,000 cells plated per well in a 96-well plate (corning, CellBind) in Opti-MEM with 10% FBS without antibiotics. Cells were immediately inoculated with BCG-BFP (MOI12). After 28 hours (day 1 after plating), 10 µg gentamicin were added per well to inactivate extracellular bacteria and cells further incubated. At the analysis day (early day 5, late day 10), supernatant was removed, cells were washed with PBS and lyzed in 0.05% SDS in PBS. Cell lysates were plated in serial dilutions and CFU counts evaluated. For the late time point at day 10, we changed the medium at day 5 and added gentamicin again. We also added BCG to wells without cells to evaluate the efficiency of bacterial killing by gentamicin. This value was used as background control (dashed line).

### Phagocytosis Assay

The omentum was isolated from β-Actin-dsred mice and digested as described above. For isolation of SC the cell suspension was stained with PDPN AF647 and CD45/CD31 FITC. Stromal cells were gated as dsred^+^ PDPN^+^ CD45/CD31^-^ cells. 12,500-15,000 SC were seeded in 2 ml Opti-Mem with 10% FBS and 0.5% Ciprofloxacin in a 6-well plate (Greiner) with and without BCG-BFP (90,000 CFU/ml). At day 3 post culture, LPM were sorted from WT mice as described above. The medium of stimulated and unstimulated SC was replaced by complete Opti-MEM supplemented with M-CSF (50 ng/ml) and 100,000 LPM were added partly. The established cultures (SC monoculture infected and uninfected and SC/LPM co-cultures infected and uninfected) were incubated for further 2 days. Afterwards, cells were harvested by scraping and a FACS staining for CD45 FITC and PDPN was performed and the percentage of BFP positive cells in the different conditions compared. Unstimulated cells served as background control. LPM were identified as CD45^+^, SC as PDPN^+^ dsred^+^ CD45^-^.

### Confocal imaging of co-cultures

For imaging in µ-Slide 8 Well high (ibidi, 80806) 5,000 SC from β-Actin-dsred mice were seeded in 200 µl complete Opti-MEM (10% FBS, 0.5% Ciprofloxacin) per well and partly stimulated with BCG-BFP (300,000 CFU/ml). At day 3, 30,000 LPM isolated from WT mice were added to some conditions and the medium was changed and replaced by complete Opti-MEM supplemented with M-CSF (50 ng/ml). At day 5, the supernatant was removed and cells washed in ice-cold PBS, followed by a 10-minute incubation in 4% PFA at room temperature in the dark. PFA was removed and the cells washed in PBS several times. Cells were stained with F4/80 AF488 1:100 for 30 minutes at room temperature in the dark slightly shaking. After washing, the cells were mounted in DAKO mounting medium and kept in the dark at 4°C till imaging. For confocal imaging of LifeAct-mice, 50,000 LPM isolated from β-Actin-dsred mice at day 3. Those were stained with PDPN AF647 in addition.

### Progenitor-SC co-cultures

SC and iMoP were isolated from WT mice as described above. 15,000 iMoP were cultured in complete Opti-MEM in 48-well plates (Corning, CellBind) with and without the addition of 5,000 SC, ± M-CSF (50 ng/ml) and ± BCG-BFP (MOI 8, 120,000 CFU/well). After approx. 24 hours, cells were harvested by scraping and FACS staining performed for Zombie Green, F4/80 PE, CD45 PerCP-Cy5.5 and CD11b APCeFL780.

### Confocal life cell imaging

Omentum was digested as described above and SC isolated from β-Actin-GFP, LifeAct-GFP or β-Actin-dsred mice and 5,000 cells plated in a µ-Slide 8 Well high (ibidi, 80806) in complete Opti-MEM and stimulated with BCG (MOI12). After 3 days, LPM were sorted from either β-Actin-GFP if SC were dsred-positive or vice versa and 50,000 cells added to the SC cultures per well. The medium was carefully exchanged to add LPM, supplemented with M-CSF (50 ng/ml) and 25 mM HEPES. The LCI chamber at LSM710 was heated before and cells were put into the incubation chamber at 37°C and 5% CO_2_. The microscope settings were adjusted quickly after addition of LPM and positions chosen in which SC had taken up bacteria to visualize how infected SC interact with Mφ.

### Retinoic acid treatment in vitro and TNF assay

SC were sorted from omentum and LPM from peritoneal lavage from WT mice as described above. 10,000 SC were seeded together with 40,000 LPM in 2 ml complete Opti-MEM with M-CSF (50 ng/ml). The co-cultures were additionally supplemented with 1.5 µg/ml retinoic acid. Control conditions received the same volume of the vector DMSO. Co-cultures were incubated for 42 hours at 37°C. Golgi-Stop (0.67 µl/ml) was added to all conditions and co-cultures partly infected with BCG-BFP (150,000 CFU/ml). Uninfected cells served as background control. After 6 hours cells were scraped and a FACS surface staining performed with PDPN AF647 and CD45 PerCPCy5.5, followed by an intracellular staining for TNF PE as described in the section FACS staining.

### Transmission electron microscopy

SC were sorted from omentum of WT mice as described and cultured in Opti-MEM in 12 well plates. SC were stimulated with BCG BFP (MOI12). In preparation for transmission electron microscopy (TEM), cultured stromal cells were fixed in a mix of 4% paraformaldehyde, 2% glutaraldehyde, and 0.1M cacodylate buffer (pH 7.4) at 4°C until contrastation. Fixed cells were then washed 3 times in 0.1M cacodylate buffer. Contrastation (1% OsO4 in H2O; 1% uranyl acetate in H2O), dehydration (30%, 50%, 70%, 90%, 100% Ethanol series), and embedding in Durcupan resin were performed using a PELCO BioWave Pro+ laboratory microwave (Ted Pella Inc.). Ultrathin sections (70 nm) were cut on a Leica Ultracut ultramicrotome (Leica microsystems) and collected on copper grids. Cells were imaged using a transmission electron microscope at 120kV (Talos L120C TEM, Thermo Fisher Scientific).

### Quantification of serum M-CSF concentrations

M-CSF concentrations in the serum were quantified using the LEGENDplex^TM^ Custom Mouse Panel 1174 (CLPX250807-AL-UF, Item No: 900008628), following the manufactur’s instructions. Sample acquisition was performed on a CytoFLEX S flow cytometer (Beckmann Coulter) on the day of staining. Data were analyzed using the cloud-based LEGENDplex^TM^ Data Analysis Software (qognit^TM^).

### Statistics

For statistical analysis, Prism 10.5.0 (GraphPad Software) was used. Results were considered statistically significant when p values were ≤ 0.05 (ns, not significant; *p < 0.05; **p < 0.01; ***p < 0.001; ****p < 0.0001). Individual statistical tests and the number of biologically independent samples are described in detail in the figure legends. In brief, a two-tailed unpaired Student’s t-test was used to compare the means of two groups. One- or two-way ANOVA was applied for multiple comparisons, followed by multiple comparison analysis. Statistical analysis was based on n, indicating the number of independent biological samples or mice, as described in the respective figure legends. If mice were removed from the dataset (e.g., due to failed infection), this is indicated in the figure legend. The analyses for the sequencing data was performed in R as described in detail before.

## Notes

### Competing Interest Statement

The authors have declared no competing interest.

### Summary of Updates

Author list updated; figures revised, including additional supplementary figures (Supplementary figure 1E, 2C, 3D); manuscript text updated.

