## Supplementary figures for "Crosstalk between Stromal Cells and Macrophages Shapes Host Immunity to Mycobacteria"

Supplementary Figure 1 (related to Figure 1)

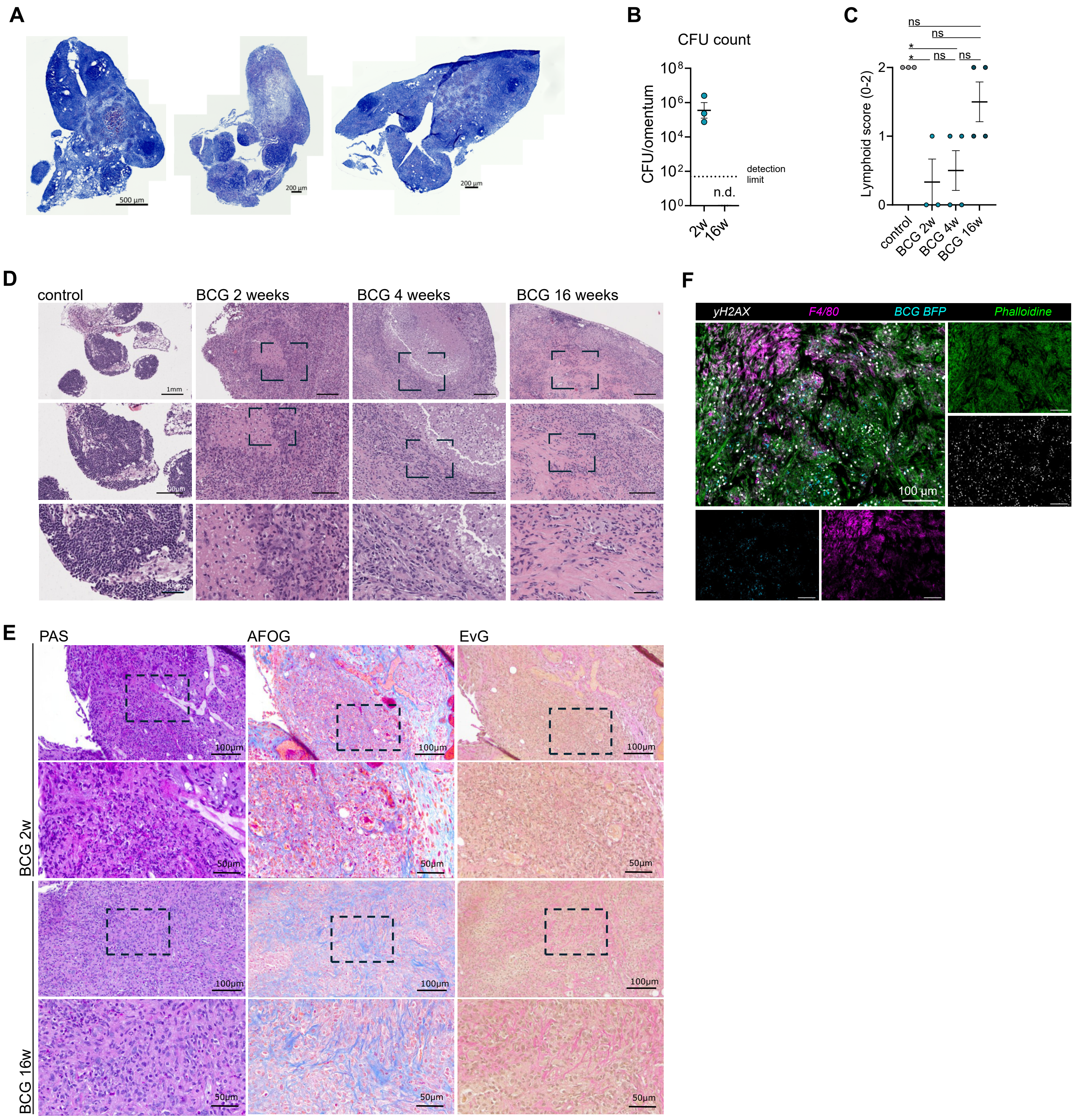

Supplementary Figure 2 (related to Figure 1)

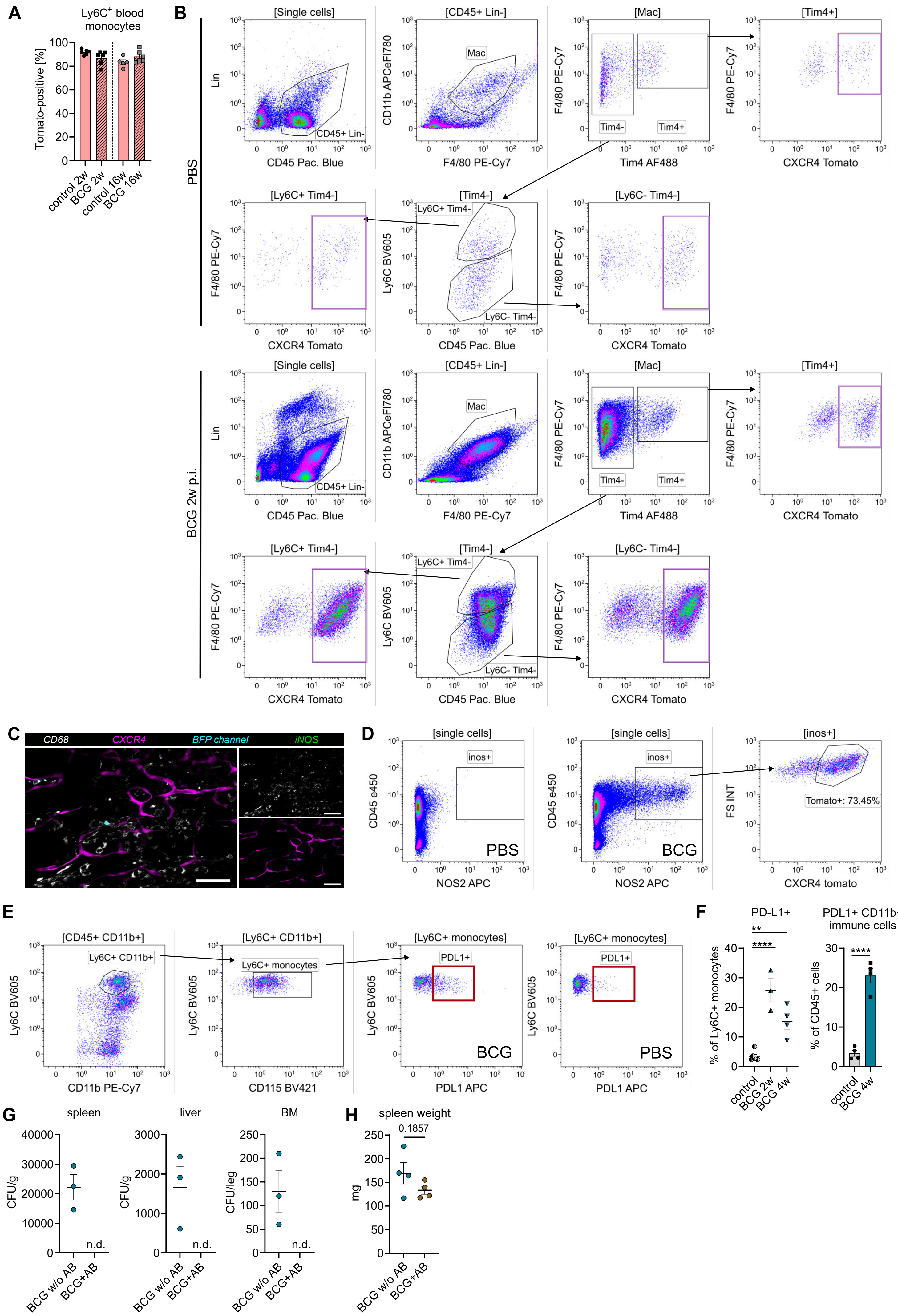

Supplementary Figure 3 (related to Figure 2)

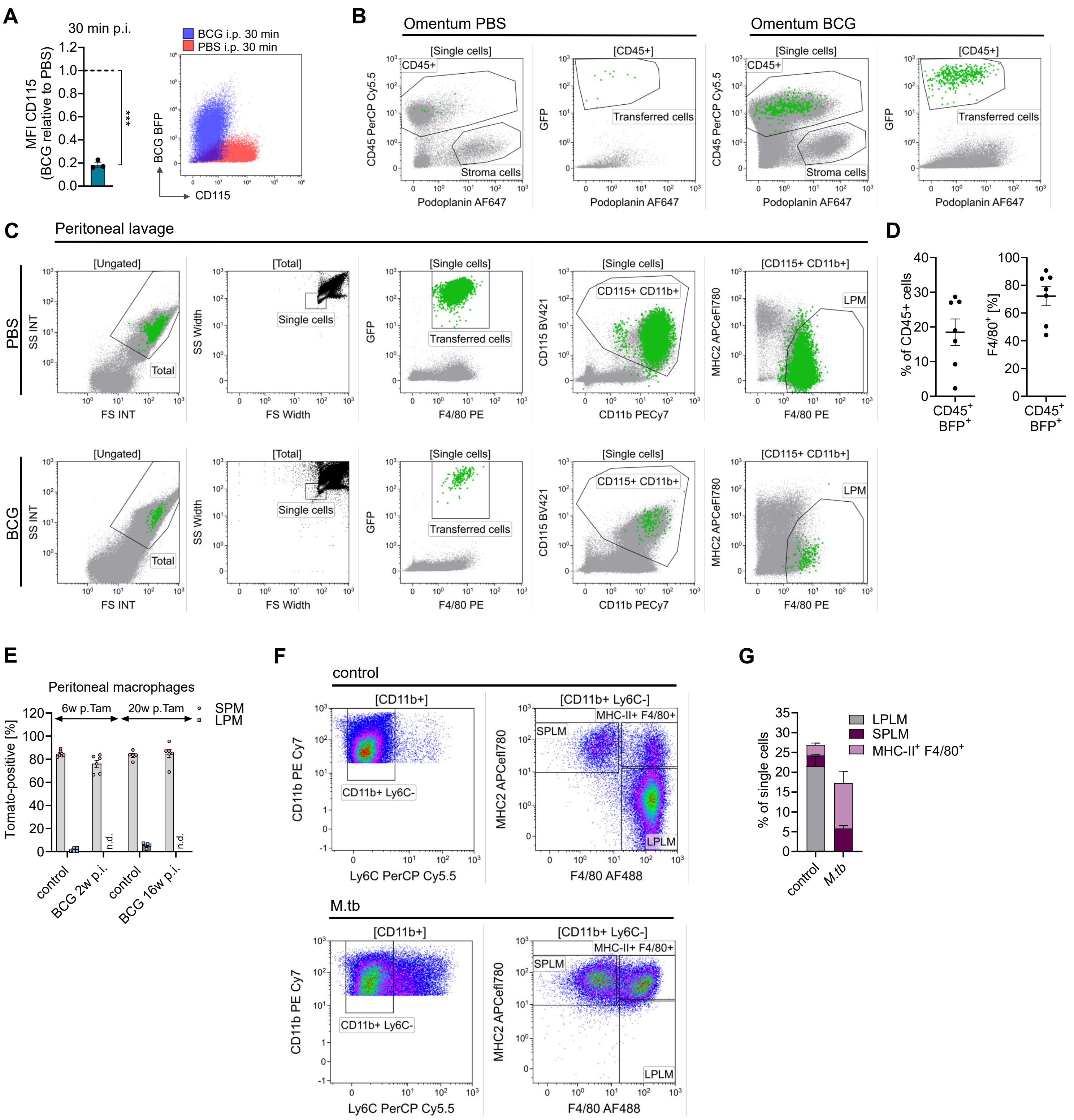

Supplementary Figure 4 (related to Figure 3)

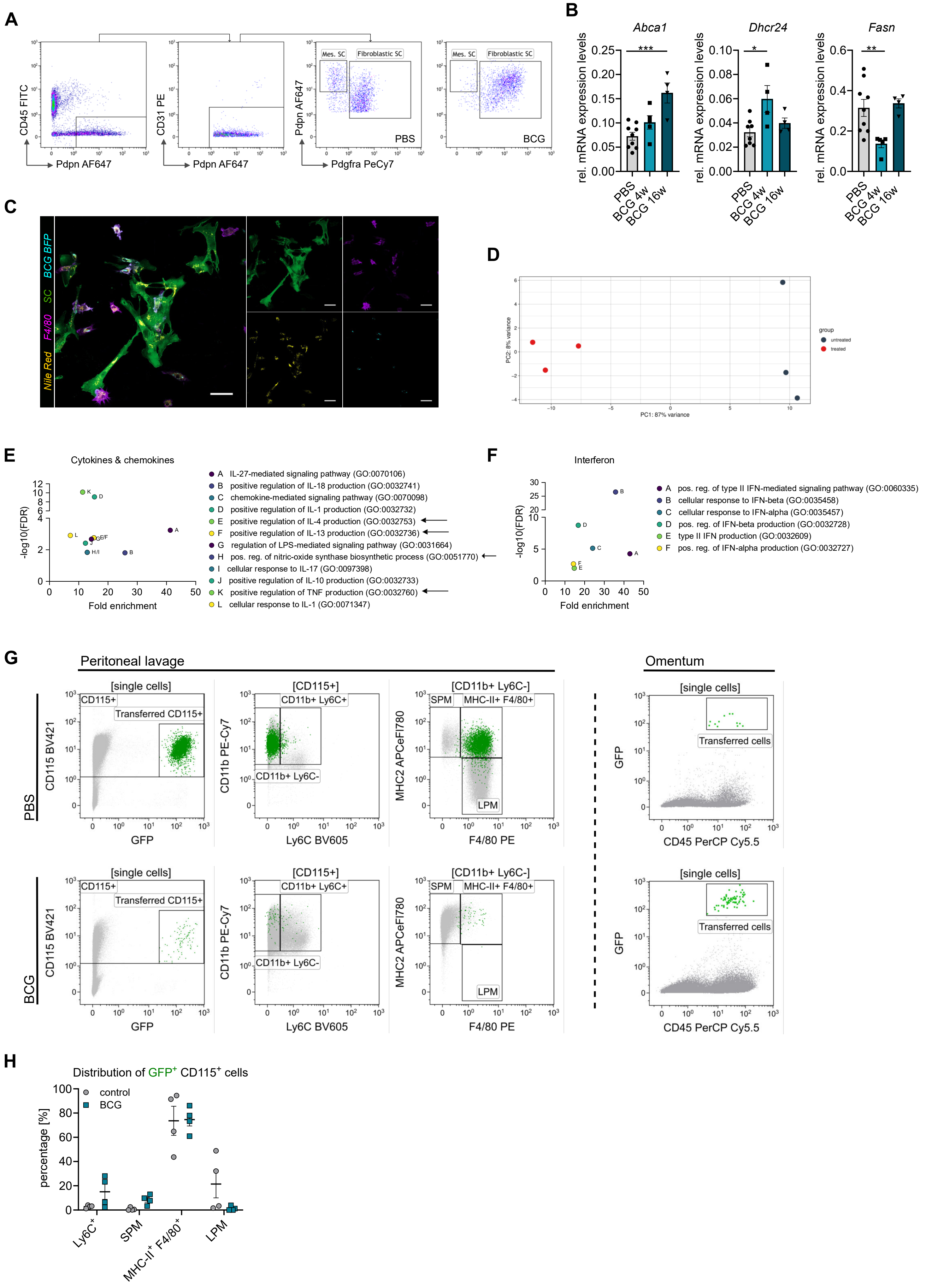

Supplementary Figure 5 (related to Figure 4)

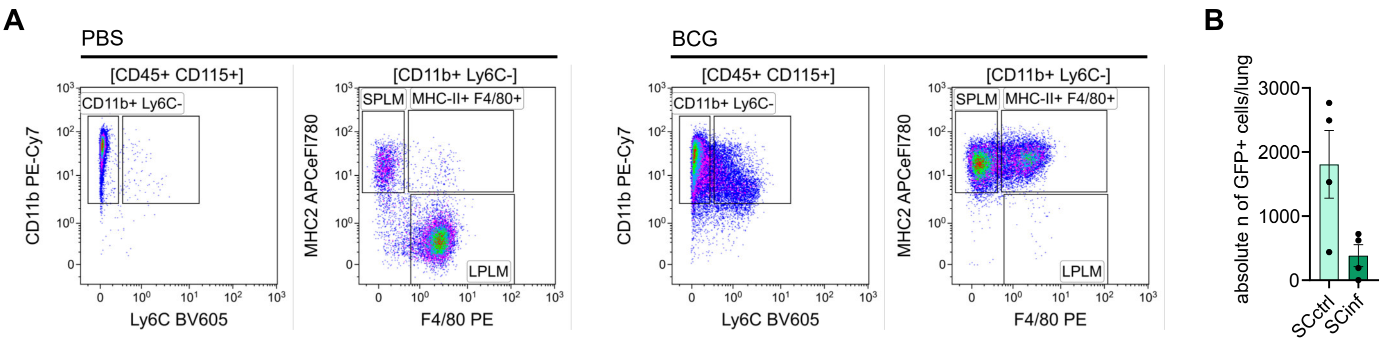

Supplementary Figure 6 (related to Figure 5)

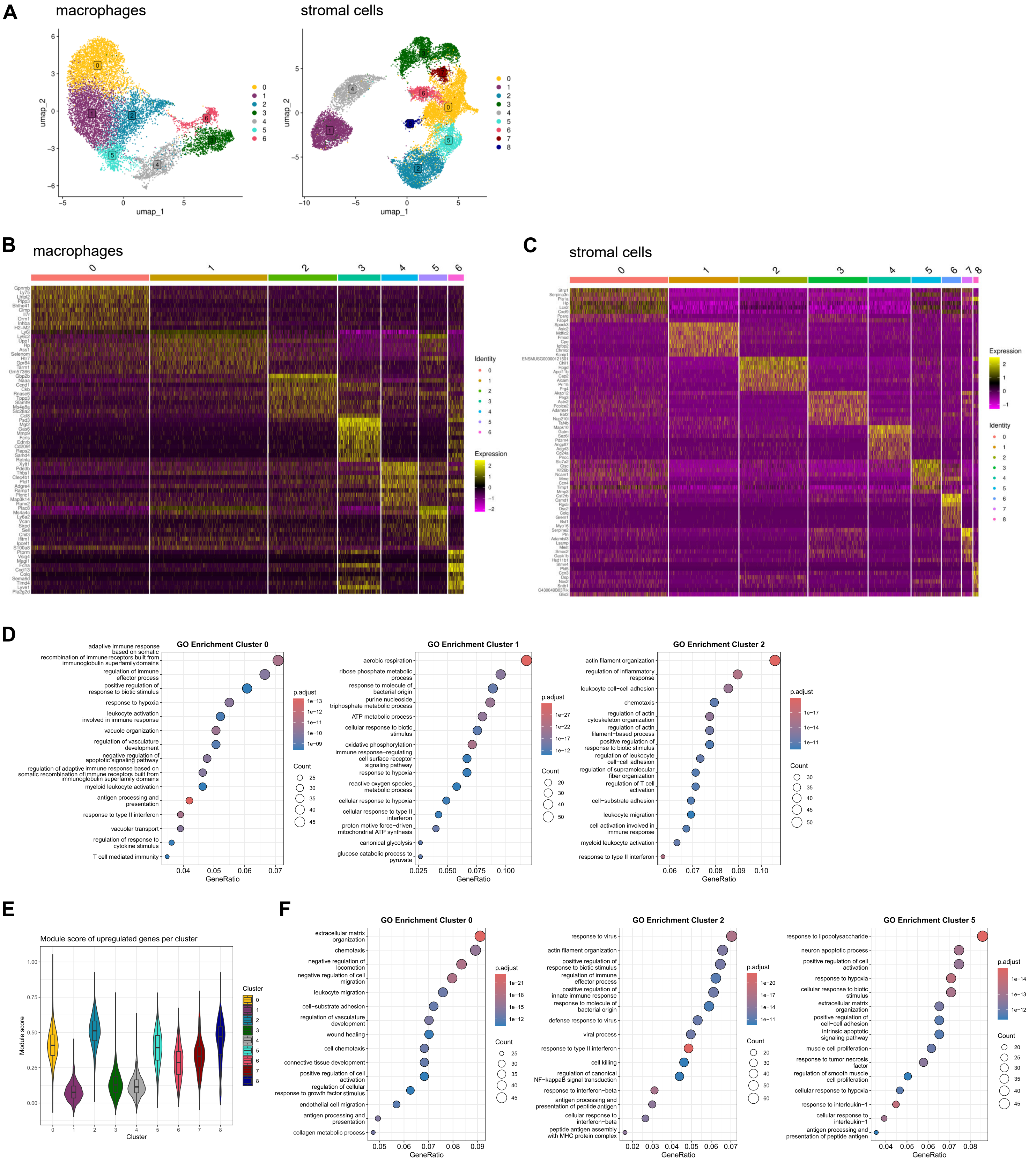

Supplementary Figure 7 (related to Figure 5)

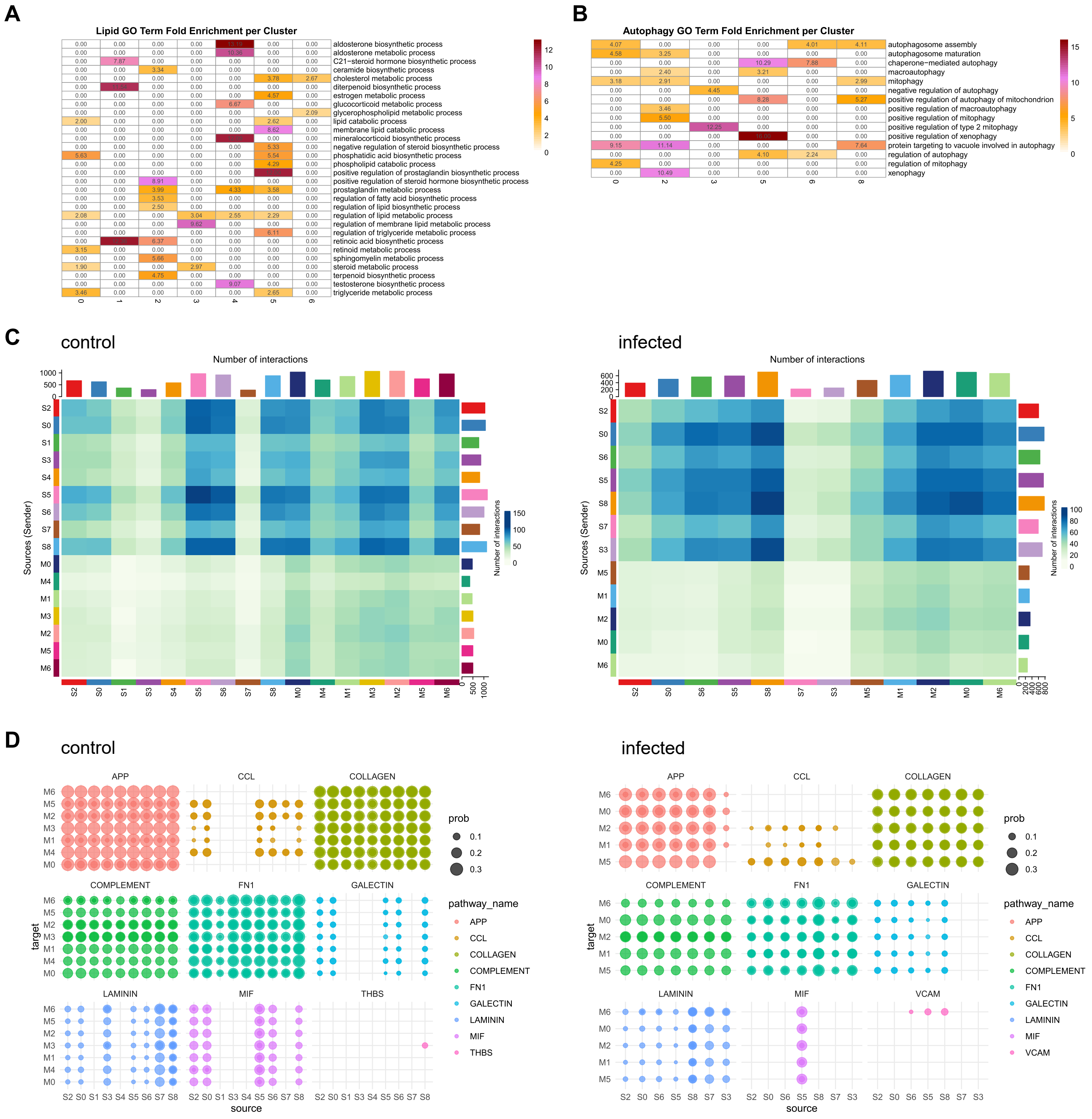

Supplementary Figure 8 (related to Figure 6)

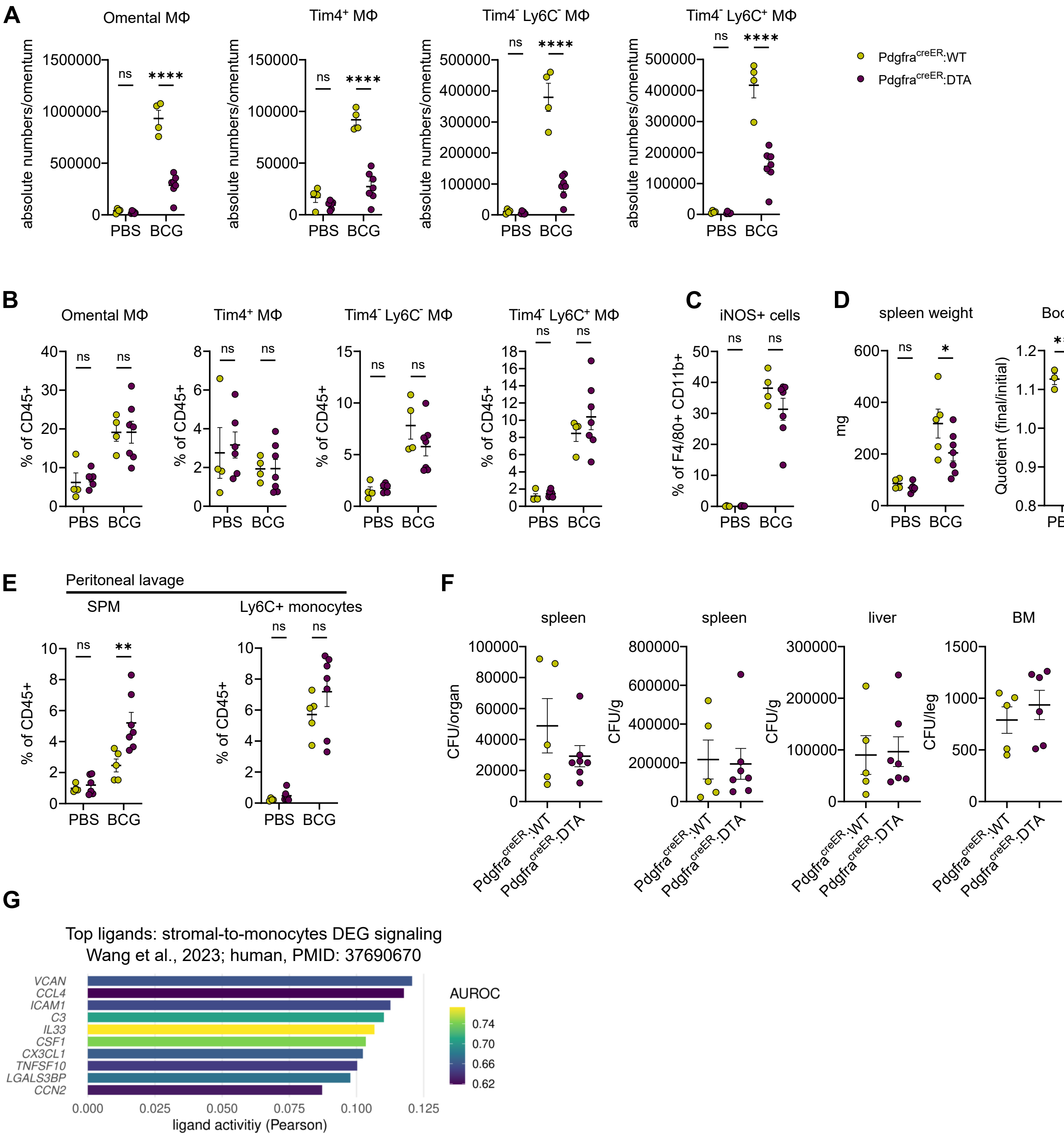
